# Microbial metabolites of flavanols in urine are associated with enhanced anti-proliferative activity in bladder cancer cells in vitro

**DOI:** 10.1101/2020.09.22.308056

**Authors:** Laura E. Griffin, Sarah E. Kohrt, Atul Rathore, Colin D. Kay, Magdalena M. Grabowska, Andrew P. Neilson

## Abstract

Dietary flavanols and their metabolites are excreted primarily via the urine, suggesting uroepithelial cells as a site of activity due to lengthy exposure to high concentrations of these compounds. Flavanols are metabolized by the gut microbiota to numerous bioavailable metabolites. The observed effects of flavanols, including cancer chemoprevention, may be due in part to the activities of microbial metabolites. Most *in vitro* mechanistic work in this area relies on a limited pool of commercially available or synthesized flavanol microbial metabolites, and little work has been done in the area of bladder cancer. The impact of physiologically relevant mixtures of native flavanols and their metabolites generated *in vivo* remains unknown. Rats were fed various flavanols after which 48 h urine samples, approximating the total bioavailable metabolome, were collected. Urine samples were profiled by UPLC-MS/MS, and their anti-proliferative activities were assayed *in vitro* in four bladder cancer cell models. Significant interindividual variability was observed for chemical profiles and anti-proliferative activities. Concentrations of microbial metabolites (valerolactones, phenylalkyl acids and hippuric acids) were positively associated with reduced bladder cancer cell proliferation *in vitro*, while native flavanols were poorly correlated with activity. These results suggest that microbial metabolites may be the primary compounds responsible for chemoprevention in uroepithelial cell following flavanol consumption. Furthermore, this highlights the potential for exploiting knowledge about individual genetics, microbiome profiles, flavonoid metabolism profiles, tumor characteristics, etc. to design personalized dietary interventions for cancer prevention and/or adjuvant therapy to reduce bladder cancer incidence and improve outcomes.

## INTRODUCTION

Flavanols (**Figure 1**) are secondary plant metabolites found in a variety of common foods such as tea, coffee, wine, chocolate, fruits, and vegetables(Cassidy & Minihane 2017). Despite their dietary abundance, flavanol bioavailability is relatively poor(Crozier et al. 2010). Flavanol monomers, such as (+)-catechin (C) and (-)-epicatechin (EC), are moderately well absorbed in the small intestine. However, oligomeric and polymeric flavanols (also called procyanidins, PCs) are poorly absorbed, and the majority of the ingested dose reaches the colon. Here, they interact with the gut microbiome, which is comprised of 100 trillion cells of high inter- and intraspecies variability(Espín et al. 2017; Behr et al. 2017). These microbes metabolize flavanols via reactions including hydrolysis, ring cleavage, and dehydration(Selma et al. 2009; Behr et al. 2017). Flavanol metabolites (representative structures in **Figure 1**) are readily absorbed into circulation and are excreted in the urine(Fernandez-Millan et al. 2014; Álvarez-Cilleros et al. 2018). While research has traditionally focused on the bioactivities of native flavanols in foods, the microbial metabolites may possess significant bioactivities as well(Monagas et al. 2010; Fernandez-Millan et al. 2014; Esposito et al. 2015; Bitner et al. 2018). The potential for flavanol microbial metabolites to confer benefits for the host is being explored(Aura 2008). Cell culture studies indicate that purified flavanol microbial metabolites exert positive effects on pathways related to glucose homeostasis and insulin signaling(Fernandez-Millan et al. 2014; Álvarez-Cilleros et al. 2018; Bitner et al. 2018). In some cases, flavanol microbial metabolites were more potent at enhancing these pathways compared to native flavanols, such as EC(Bitner et al. 2018). Interestingly, Esposito et al.(Esposito et al. 2015) reported that the anti-obesity benefits of black current anthocyanins (flavonoids similar to flavanols) in mice were lost when antibiotics (Abx) were simultaneously administered, suggesting that the microbiome is critical for mediating the benefits of flavonoids (potentially via bioavailable microbial metabolites). This growing body of evidence suggests that flavanol microbial metabolites may play a significant role in achieving the observed benefits of flavanol consumption, given their potential bioactivity and the fact that they are generally more bioavailable than the native forms. However, the majority of the *in vitro* mechanistic work in this area has relied on individual purified flavanol microbial metabolite compounds. The impact of physiologically relevant mixtures of native flavanols and metabolites generated *in vivo* remains unknown.

**Figure 1.**
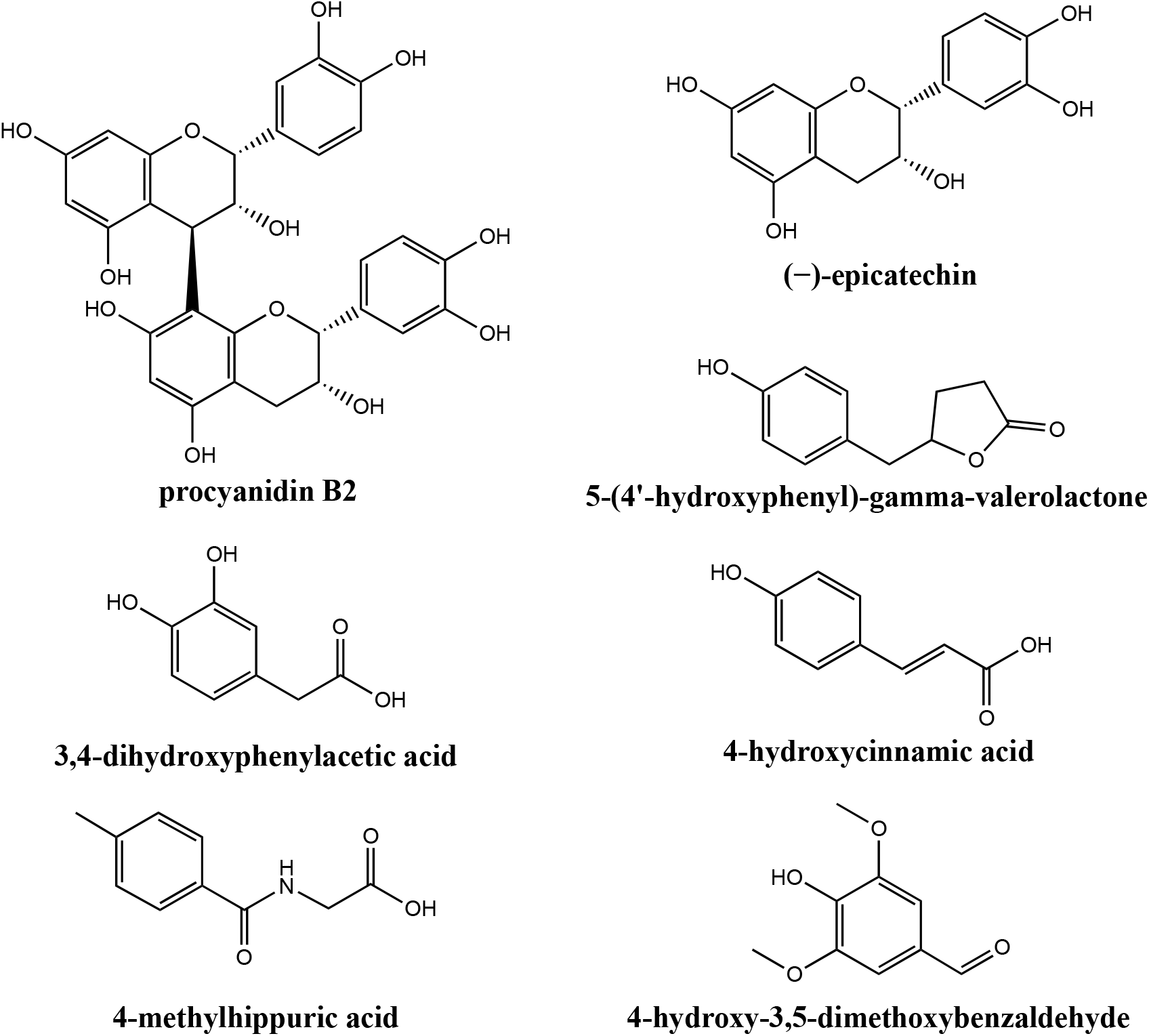
Structures of representative flavanols [procyanidin B2 and (−)-epicatechin] and representative flavanol microbial metabolites.

It is well known that absorbed flavanols and their microbial metabolites are rapidly metabolized by Phase-II detoxification systems, which enhances their water solubility and thus excretion into the urine for elimination(Williamson et al. 2018). This is generally thought to reduce the potential impacts of bioavailable flavanols in peripheral tissues. However, this presents a unique opportunity where flavanols and their microbial metabolites are exposed to uroepithelial cells (particularly the bladder) at high concentrations for extended times. This suggests the potential for chemoprevention of bladder diseases via dietary flavanol administration. Bladder cancer is the fourth leading cause of cancer in men in the US, with lower incidence in women(Siegel et al. 2017). In 2020, bladder cancer will contribute to an estimated 18,000 deaths(Siegel et al. 2017). Bladder cancer is diagnosed as non-muscle invasive bladder cancer, the more common and relatively non-lethal but recurring diagnosis, and muscle-invasive bladder cancer, an aggressive and potentially metastatic diagnosis(McConkey & Choi 2018).

Prevention, as well as complementary strategies to optimize traditional therapy, are needed to reduce the incidence and improve outcomes of bladder cancer. Bladder cancer therapy largely depends on invasiveness and tumor stage. For non-muscle invasive disease, aggressive monitoring via cystoscopy and removal of recurring lesions via Transurethral Resection of Bladder Tumor are common, along with Bacillus Calmette-Guerin treatment(Chang et al. 2016). In these patients, decreasing the rate of recurrence is needed. For muscle-invasive disease, treatment is more aggressive, including combinatorial chemotherapy and radiation regimens or cystectomy with or without neoadjuvant chemotherapy(Chang et al. 2016). As bladder cancer is largely a disease of older patients, neoadjuvant chemotherapy is not always feasible. In order to attain some level of tumor downstaging prior to cystectomy in this frail patient population, complementary therapies have long been sought. Therefore, phytochemicals such as flavanols could play a role in improving bladder cancer incidence and survival. Epidemiological data suggest that consumption of flavanols may impact bladder cancer development and recurrence. In the European Prospective Investigation into Cancer and Nutrition (EPIC) study, flavanol intake inversely correlated with bladder cancer risk(Zamora-Ros et al. 2013). Non-smoking bladder cancer patients with a high consumption of green tea (which contains flavanols such as epigallocatechin-3-gallate, EGCG) were less likely to undergo bladder cancer recurrence, up-staging, and up-grading(Furukawa et al. 2017). Importantly, EGCG accumulates in the normal bladder and bladder tumors(Gee et al. 2017). *In vitro*, the bulk of flavanol studies have focused on EGCG treatments of cell lines and demonstrated antiproliferative effects in bladder cancer cell lines(Philips et al. 2009; Luo et al. 2017). However, these studies did not include flavanol microbial metabolites, which may be more potent than their native precursors due to greater circulating levels.

The purposes of this study were to 1) determine the impact of the gut microbiome on the profile of systemically bioavailable compounds (native and microbial metabolites) and 2) determine the contribution of the gut microbiome to the bioactivity of circulating species using *in vitro* models of bladder cancer, following consumption of comparatively bioavailable (C/EC) and poorly-bioavailable (PCs) flavanols. We hypothesized that flavanol microbial metabolites will be present in urine following dietary flavanol exposure and that these metabolites will be abolished by antibiotic (Abx) treatment. Further, we hypothesized that urine from animals fed flavanols would have significant antiproliferative effects on *in vitro* models of bladder cancer and that these activities would be significantly reduced when the gut microbiome was diminished due to Abx treatment. We also hypothesized that the reduction in activities due to Abx would be greater for poorly-bioavailable (PCs) vs. comparatively bioavailable (C/EC) flavanols.

## METHODS

### Flavanol standards and extracts

Representative flavanol monomers, (+)-catechin hydrate (C) and (-)-epicatechin (EC), were purchased from Millipore Sigma (Burlington, MA). Vitaflavan^®^ grape seed extract (GSE), a flavanol extract rich in oligomeric flavanols (PCs) was purchased from DRT Nutraceutics (Dax, France). Manufacturer specifications for Vitaflavan can be found in **Table 1**. Prior to oral gavage procedures, suspensions of 1:1 C/EC (125 mg/mL) or GSE (125 mg/mL) were prepared in water. Mixtures were sonicated for 5 min prior to use.

**Table 1.** Composition of Vitaflavan^®1^.

|  |  |
| --- | --- |
| Total Polyphenol Content | >96% |
| Monomers | <25% |
| Dimers + Trimers | >30% |
| Procyanidin Content (GPC) | >75% |
| Procyanidins Content (Porter) | 70 |

<sup>1</sup>Information provided by DRT Nutraceuticals

### Animals and diets

The Institutional Care and Use Committee at the David H. Murdock Research Institute approved all animal procedures (protocol #19-005). Male outbred Wistar rats (n = 30, 6 weeks old, ~225 g) were purchased from Charles River (Wilmington, MA). Rats were individually housed under standard conditions (12 hr light/dark cycle, 30-70% relative humidity, 20-26°C). Food and water were provided *ad libitum*. Upon receipt, rats were acclimatized to laboratory conditions on standard chow diet for 5 days prior to the start of the experiment. Water was provided in standard bottles. After acclimatization, rats were transitioned to the purified AIN-93G Growing Rodent Diet (D10012G) from Research Diets Inc. (New Brunswick, NJ) and randomized to one of six treatments (**Table 2**), *n* = 5 rats/treatment.

**Table 2.** Experimental treatment groups.

| **Treatment** | **Antibiotics** | **Treatment Name** | **Treatment Abbreviation** |
| --- | --- | --- | --- |
| water | – | Vehicle | V |
| 1:1 C/EC | – | Monomer Control | M |
| GSE | – | GSE control | GSE |
| water | + | Antibiotic Control | V + Abx |
| 1:1 C/EC | + | Monomer + Antibiotics | M + Abx |
| GSE | + | GSE + Antibiotics | GSE + Abx |

### Antibiotic treatment

The rats assigned to Abx treatment groups (V/A, M/A, GSE/A) received a cocktail consisting of 0.5 g/L vancomycin (Triangle Compounding Pharmacy, Cary, NC), 1 g/L neomycin sulfate (Huvepharma, Maxton, NC), 1 g/L metronidazole (Unichem Laboratories, Hasbrouck Heights, NJ), 1 g/L ampicillin (Auromedics Pharma, East Windsor, NJ) administered in drinking water. This cocktail has the demonstrated ability to sufficiently reduce the gut microbiome in rodent models(Esposito et al. 2015). Rats were maintained on Abx for 10 days prior to oral gavage. Abx were refreshed every 48 hours.

### Flavanol metabolite collection

Rats were fasted overnight before oral gavage. Rats were gavaged in groups of *n*=6 at a time, one from each treatment group. Flavanol solutions (C/EC and GSE) were prepared fresh each morning. Urine collection tubes were pre-filled with formic acid and ascorbic acid so that the final concentrations were 0.2% formic acid and 0.01 M ascorbic acid when diluted with urine (based on an estimate of 10 mL urine/kg/d). Rats were gavaged 1 mL of either water, 125 mg/mL C/EC, or 125 mg/mL GSE via a 16-gauge curved gavage needle (Kent Scientific, Torrington, CT,). Following gavage, rats were placed individually in metabolic cages for 48 h for urine collection. Rats were allowed food and water *ad libitum* and Abx treatments were continued throughout collection. Rats were euthanized upon removal from metabolic cages. Urine samples were frozen at −80°C.

### Flavanol metabolite purification

Following the urine collection procedure for all rats, urine samples were freeze-dried until all water was removed from the samples. Once dry, the urine samples were reconstituted in 5 mL of 0.1% formic acid in 1:1 methanol:ethanol. Samples were vortexed, sonicated, and centrifuged at 10,000 x *g* to assist in the removal of urea and salts(Pinho & Macedo 2005). Supernatants were collected and sterile filtered with Millex 0.22 μm Millipore membrane filters (Sigma, Burlington, MA) into microcentrifuge tubes in 1 mL aliquots. Samples were then speedvapped with a Savant DNA 120 SpeedVac Concentrator (Thermo Fisher, Waltham, MA) to dryness for 2 hours. Dried flavanol extracts were stored at −80°C when not in use and were reconstituted in a standardized volume of water prior to usage in cell culture.

### Urine composition analysis

Analysis was performed based on our established workflow(Nieman et al. 2018; Chandra et al. 2019). To establish the metabolome assay of flavanol metabolites a systematic review approach using PubMed was utilized to capture reported composition data from published works. References standards were purchased where available or obtained from an in-house reference standard library. Assay development workflow included fragment/transition optimization via direct syringe infusion of pure reference standards. Sequential optimization of scanning and integration parameters were established in multiple reaction monitoring (MRM), scheduled MRM (sMRM) and advanced scheduled MRM (ADsMRM) modes. A mixture of reference standards was spiked in pooled extracted “blank” and injected multiple times at concentrations reflecting lower limits of quantification (LLOQ), middle quality control (MCQ), and upper limit of quantitation (ULOQ) to establish chromatography drift, signal stability, background noise and integration thresholds. Signals for metabolites for which no reference standards were commercially available were optimized using MRM scanning mode, where transitions were programmed for their optimized precursor base structures (i.e., unconjugated/pre-metabolized analyte) and known biological conjugate masses, such as glucuronide, sulfate, and glycine. The MRM’s were run on archived pooled urine samples from a human flavanol intervention as a series of dilutions to identify the linearity of signals (i.e., signals proportionally affected by overlapping precursor and product signals). Identification was based on fragmentation profiling of the precursor structure and 3-5 product transitions. The method was merged with the previous ADsMRM method. Finally, the pooled samples were scanned for 485 additional analytes present in our in-house metabolome database to identify previously unreported metabolites. The data from these scans was merged with the previous ADsMRM method as described above. This final targeted/broad-spectrum quantitative assay was optimized and validated to detect 108 analytes, which were quantified relative to 94 authentic commercial and synthetic standards. Where reference standards for metabolites (including Phase II conjugates) were not available (14 analytes), identification was based on fragmentation profiling involving the precursor structure and 3–5 product transitions. These metabolites were quantified relative to their closest structural reference standard with similar ionization intensities (the MRM table for all analytes reported in this study is shown in **Supplementary Table 1**).

Analytes were purified from 50 μL rat urine by 96-well solid phase extraction (Strata™-X Polymeric Reversed Phase, microelution 2 mg/well). Precision was measured based on %CV of area ratios and accuracy was evaluated with %CV of calculated concentration and retention time at three different concentrations (near LLOQ, MCQ, and ULOQ). The solid phase extraction treated samples were chromatographically separated and quantified using Exion ultra-high-performance liquid chromatography-tandem mass spectrometry (SCIEX^®^ QTRAP 6500^+^) with electrospray IonDrive Turbo-V Source. The samples were injected into a Kinetex PFP UPLC column (1.7μm, 100Å, 100mm x 2.1mm; Phenomenex^®^) with oven temperature at 37°C. Mobile phase A and B consisted of 0.1% v.v. formic acid in water and 0.1% v.v. formic acid in LC-MS grade acetonitrile, respectively, with a binary gradient ranging from 2% to 90% B over 30min and flow rate gradient from 0.55 mL/min to 0.75 mL/min. MS/MS scanning was accomplished by ADsMRM using polarity switching between positive and negative ionization mode in Analyst (v.1.6.3, SCIEX), with quantitation using MultiQuant (v.3.0.2, SCIEX) software platforms. Internal standards included L-tyrosine-^13^C_9_,^15^N, resveratrol-(4-hydroxyphenyl-^13^C_6_), phlorizin dehydrate (Sigma-Aldrich), theobromine-d_6_, caffeine-^13^C_3_, hippuric acid-^13^C_6_, and propyl paraben-^13^C_6_ (Toronto Research Chemical, TRC; Toronto, Canada).

Calibration curves containing 11 points (0.001-10 μM) to 14 points (0.001-100 μM) were established by spiking reference standards in matrix matched SPE extracted Surine (synthetic urine negative control; S-020, Sigma-Aldrich Corporation, St. Louis, MO). Source parameters included: curtain gas 35, ion-spray voltage 4500 V, temperature 550 °C, nebulizer gas 70, and heater gas 70. The optimized analyte specific quadrupole voltages (mean ± SD) for negative mode were 53 ± 28 V for declustering potential, 10 ± 1 V for entrance potential, 28 ± 10 V for collision energy, and 12 ± 6 V for collision cell exit potential. Similarly, for positive mode the optimized analyte specific quadrupole voltages (mean ± SD) were 46 ± 21 V for declustering potential, 10 ± 1 V for entrance potential, 22 ± 10 V for collision energy, and 15 ± 5 V for collision cell exit potential.

### Cell Culture

SCaBER (HTB-3), RT4 (HTB-2), SW780 (CRL-2169), and HT-1376 (CRL-1472) cells were purchased from ATCC. These cell lines were chosen in order to represent two luminal cell lines (RT4, SW780) and two basal cell lines (SCaBer, HT-1376)(Warrick et al. 2016) and represent female (SW780, HT-1376) and male (SCaBer, RT4)-derived tumor cell lines(Rasheed et al. 1977; Zuiverloon et al. 2018). Cells were grown in 96 well plates at a range of seeding densities to optimize a final density yielding 80-90% confluency after four days. SCaBER cells were seeded at 5,000 cells/well in MEM alpha + nucleosides medium with 10% FBS. RT4 cells were seeded at 20,000 cells/well in McCoy’s medium with 10% FBS. SW780 cells were seeded at 5,000cells/well in RPMI-1640 + L-glutamine media with 10% FBS. HT-1376 cells were seeded at 10,000 cells/well in MEM alpha + nucleosides media with 10% FBS. Cells were plated then incubated overnight before adding the urine samples. To make a 10% final concentration of urine in media, 80uL of each sample was added to 720 *μ*L of media. 100 *μ*L of each 10% sample in media was added per well of a 96 well plate. Six technical replicates were used for each sample. Blank wells and water-treated wells were used as controls. Cell growth was assessed using a modified crystal violet assay(Kohrt et al. 2020). In brief, cells were fixed using 4% paraformaldehyde in PBS (Alfa Aesar) and stained with 0.05% crystal violet (Ricca). Stain was collected using 10% acetic acid and quantified in a spectrophotometer at 590 nm (SpectraMax ID3, Molecular Devices). The average stain quantification was generated from the water control group and each experimental value was divided by this average to produce a normalized fold change value. These values were expressed as % of the water control.

### Data analysis and statistics

For cell culture experiments, individual data points are reported as the means from *n*=6 wells/individual rat urine sample. The cell culture experiment was replicated, and results of both experiments were generally in agreement. Analyses were performed to determine predictive relationships between urine composition and inhibition of bladder cancer growth. Due to the extensive list of analytes and multiple cell lines, individual compounds were not analyzed for correlations with activity. Associations between groups of compounds and inhibitory activities were thus performed for groups of compounds in order to establish general relationships between compound classes and activity. The data points were the calculated composition values for each urine sample, and the mean % proliferation from n=6 replicate wells for each urine sample (*n*=5 48-h urine samples, one per rat, per treatment) and cell line. Data and statistical analyses were performed on Prism 8.4.2 software (GraphPad, La Jolla, CA).

For concentrations of analytes total in urine samples, 2-way ANOVA was performed to determine the statistical significance of main effects (flavanol and Abx treatment) and interactions. If a significant main effect or interaction was detected, Holm-Sidak *post hoc* tests to account for multiple comparisons were performed to determine differences among the 3 flavanol treatments within each antibiotic treatment group (Control and Abx). Holm-Sidak post hoc tests were also performed to determine differences between antibiotic treatments (Control and Abx) for each flavanol. The overall family-wise error rate was set as 0.05, with one family per group.

For proliferation in various bladder cancer cell lines (expressed as a % relative to control), 2-way ANOVA was performed to determine the statistical significance of main effects (flavanol and Abx treatment) and interactions. If a significant main effect or interaction was detected, Holm-Sidak *post hoc* tests to account for multiple comparisons were performed to determine differences among the 3 flavanol treatments within each antibiotic treatment group (Control and Abx). Holm-Sidak *post hoc* tests were also performed to determine differences between antibiotic treatments (Control and Abx) for each flavanol. The overall family-wise error rate was set as 0.05, with one family per group.

Proliferation was compared between the highest vs. lowest quartiles (lowest vs. highest % proliferation compared to control) for each individual urine samples in each bladder cancer cell line. Proliferation was compared between the highest vs. lowest quartiles by t-tests, with the Holm-Sidak method to control for multiple comparisons (1 family, 4 t-tests: 1 per cell line), with the overall family α= 0.05. Each cell line was analyzed individually, without assuming a consistent SD.

Within each cell line, concentrations of each analyte class were compared between the highest vs. lowest quartiles by t-tests, with the Holm-Sidak method to control for multiple comparisons (1 family per cell line, 6 t-tests: 1 per compound class).

Within each analyte class, proliferation was compared between bladder cancer cells exposed to the highest vs. lowest quartiles of analyte concentration by t-tests, with the Holm-Sidak method to control for multiple comparisons (1 family per compound class, 4 t-tests: 1 per cell line).

See figure captions for complete descriptions of each analysis. For all analyses, α was set *a priori* at 0.05. Data are presented as mean ± SEM.

## RESULTS AND DISCUSSION

### Urine composition

Analytes from the constructed method that were determined *post hoc* to be not applicable to the administered treatments, and those exhibiting poor linearity for the external standards, were excluded from the analysis. Signals with intensities at or below the lower limit of detection were reported as 0 μM. Peaks with intensities between the lower limits of detection and quantification were reported as 0.0001 μM. Based on these criteria, 75 analytes (from the original 108 measured analytes) were selected for further analysis due to their relevance to the substrates fed to the animals, or their presence at detectable levels in the samples. These 75 compounds detected in urine were then classified categorically based on the three categories: 1) native flavanol or flavanol microbial metabolites, 2) unconjugated form or Phase-II conjugated, and 3) metabolite stage: native flavanol, valerolactone, phenylalkyl acid or related, cinnamic acids or related, benzoic acids or related, hippuric acids or related, other aromatics, non-aromatics. The categorizations for each compound are shown in **Supplementary Table 2**. Individual compound levels are presented in **Supplementary Table 3**. Urine composition as a function of treatment was determined for total native flavanols, flavanol microbial metabolites, valerolactones, phenylalkyl acids, cinnamic acids, hippuric acids, non-aromatics, other aromatics and benzoic acids, and the results are presented in **Figure 2**.

**Figure 2.**
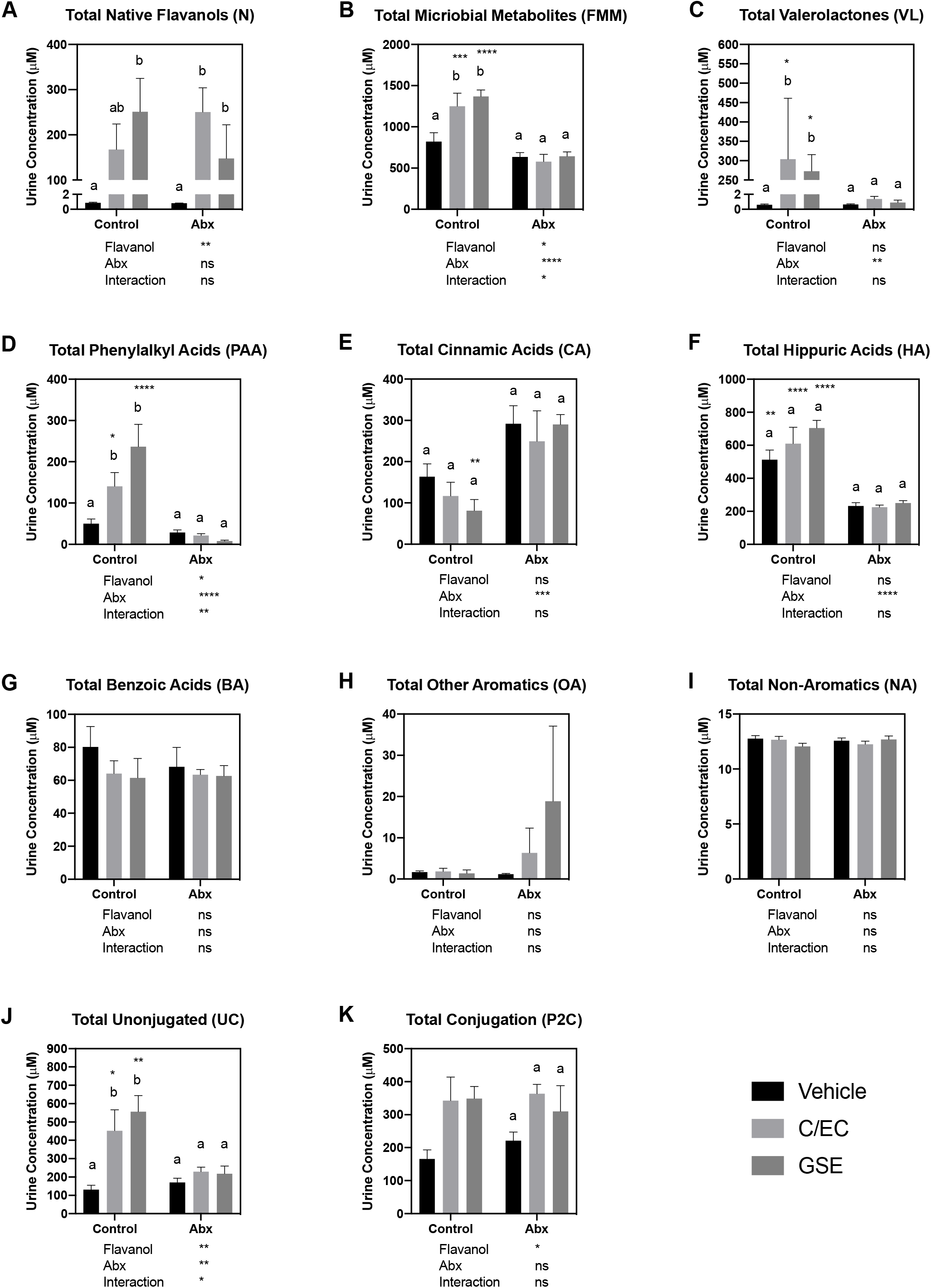
Concentrations of total levels of native flavanols, flavanol microbial metabolites, and their phase-II conjugates in urine samples. Values are presented as mean ± SEM. For each measure, 2-way ANOVA was performed to determine the statistical significance of main effects (flavanol and Abx treatment) and interactions. Values below each graph indicate the results of 2-way ANOVA. If a significant main effect or interaction was detected, Holm-Sidak post hoc tests to account for multiple comparisons were performed to determine differences among the 3 flavanol treatments within each antibiotic treatment group (Control and Abx); bars not sharing a common superscript letter within each group are statistically different. Holm-Sidak post hoc tests were also performed to determine differences between antibiotic treatments (Control and Abx) for each flavanol; asterisks indicate a significant difference for that flavanol between Control and Abx (\**P*<0.05, \*\**P*<0.01, \*\*\**P*<0.001, \*\*\*\**P*<0.0001). The overall family-wise error rate was set as 0.05, with one family per group.

For total native flavanols (**Figure 2A**), the administered flavanol (control, C/EC or GSE) significantly affected the total levels detected, but Abx administration did not. Levels of native flavanols in V treatments were minimal, and levels in the C/EC and GSE urines were on the order of 100-300 μM. However, there was significant interindividual variation in total native flavanols detected for both M and GSE treatments, which resulted in few statistically significant differences between treatments. It is worth noting that due to complexities associated with measuring procyanidins, our method detected C, EC, PC B1, PC B2 and PC C1 and their phase-II conjugates; other PCs present in GSE were not detected. These results suggest that the overall native flavanol bioavailability is not significantly affected by Abx administration (and loss of the microbiome).

For total flavanol microbial metabolites (**Figure 2B**), both the nature of the flavanol and the administration of Abx significantly influenced total levels of metabolites detected. Both C/EC and GSE significantly elevated total flavanol microbial metabolites by roughly 2-fold compared to vehicle, and Abx administration effectively prevented these changes. These data demonstrate that 1) our Abx effectively eliminated microbial metabolism of flavanols, 2) flavanol microbial metabolite production varies by the nature of the flavanol substrate, and 3) significant overlap is observed between flavanol microbial metabolites and some the same compounds from different sources (existing naturally in the diet, or microbial derived from non-flavanol precursors in the diet). Due to the fact that high doses of C/EC and GSE both only doubled the detected flavanol microbial metabolite levels compared to vehicle, the contribution of these flavanol microbial metabolites from non-flavanol sources is significant.

We then examined flavanol microbial metabolite classes. For valerolactones (**Figure 2C**), the nature of the flavanol did not affect detected levels but administration of Abx significantly reduced total levels detected. Both C/EC and GSE appear to be equally effective substrates for valerolactone synthesis. Similar to native flavanols, levels of valerolactones in vehicle treatments were minimal, and levels in the M and GSE urines were on the order of 250-300 μM. However, Abx completely eliminated the presence of valerolactone down to vehicle levels, confirming that while many flavanol microbial metabolites can be derived from non-flavanol sources (as seen above), valerolactones do not come from the background diet and depend on an intact microbiome for synthesis.

For total phenylalkyl acids (**Figure 2D**), both the nature of the flavanol and Abx administration significantly affected total levels detected. Both C/EC and GSE produced large increases in phenylalkyl acids (>100 μM above vehicle control), but GSE appears to be a better substrate for phenylalkyl acid synthesis. Abx completely eliminated the presence of phenylalkyl acids down to vehicle levels, confirming that phenylalkyl acids are similar to valerolactones in that only low levels come from the background diet and they depend on an intact microbiome for synthesis.

Total cinnamic acids (**Figure 2E**) were unique among all measured flavanol microbial metabolites. The nature of the flavonoid did not affect detected levels but administration of Abx significantly increased total levels detected for the GSE treatment. These data suggest that significant cinnamic acid levels are obtained from the background purified diet (from sources such as cereal grains)(Stuper-Szablewska & Perkowski 2019), either as preformed cinnamic acids or as microbial metabolites from larger substrates. It appears that the latter case in more probable, as cinnamic acid levels were higher in Abx groups (albeit only significantly so for GSE). The trend of higher cinnamic acid levels in Abx animals could be explained by the presence of preformed cinnamic acids in the diet, which cannot be metabolized by the microbiome when Abx are administered and thus remain high.

Hippuric acids (**Figure 2F**) were by far the most abundant flavanol microbial metabolites for all groups, comprising ~50% of all detected flavanol microbial metabolites. The nature of the flavonoid did not affect detected hippuric acid levels but administration of Abx significantly reduced total levels detected. Hippuric acids are clearly derived in significant quantities from the background diet, which is due to its role as a primary end product of the metabolism of both amino acids and aromatics(Phipps et al. 1998; Pero 2010). The large reduction observed in Hippuric acid levels by Abx administration strongly support the dependency of these metabolites on the microbiome.

Concentrations of benzoic acids, other aromatics and non-aromatics (**Figure 2G-I**) were overall lower than other flavanol microbial metabolites. Benzoic acid levels were not significantly affected by the nature of the flavanol nor Abx administration. Benzoic acids appear to derive significantly from the diet as pre-formed compounds, as their formation did not depend on the intact microbiome. Other aromatic levels were also not significantly affected by the flavonoid nor Abx. However, there were elevated (albeit nonsignificant) levels of other aromatics in the GSE/Abx group, suggesting that appreciable levels of other aromatics originate from GSE but are degraded in the presence of an intact microbiome.

The previous analyses included unconjugated compounds and phase-II conjugated (*O*-methylated, glucuronidated and/or sulfated) metabolites for both native flavanols and flavanol microbial metabolites. In order to determine whether treatments had an effect on detoxification, we examined total levels of unconjugated and phase-II conjugated compounds (totals for both native flavanols and flavanol microbial metabolites). For total unconjugated compounds (**Figure 2J**), both the nature of the flavonoid and Abx administration significantly influenced overall conjugates. Interestingly, this followed the pattern observed for total flavanol microbial metabolites, valerolactones and phenylalkyl acids but not total native flavanols. This may be explained by the fact that a significant reduction in total circulating substrates in Abx animals due to decreases in flavanol microbial metabolite production, with little change in native flavanols, meant less substrate competition for phase-II enzymes, and hence greater conjugation and reduced levels of unconjugated compounds. However, this interpretation is not supported by total phase-II conjugated compounds. For phase-II conjugated compounds (**Figure 2K**), the administered flavanol significantly affected the total levels detected, but the administration of Abx did not. An alternative explanation is that unconjugated metabolites were reduced due to redcued enzyme induction during Abx administration. This is supported by the fact that total native polyphenol absorption was unaffected by Abx. However, the majority of the microbial metabolites of polyphenols were affected by Abx. This was reflected by reductions in valerolactones, phenylacetic and hippuric acids in the Abx group, which represented ~75% of the recovered metabolites. The remaining ~25% of recovered polyphenol metabolites, namely, cinnamic acids, benzoic acids and other aromatics, were not affected by reduction in the microbiome. Interestingly conjugated (Phase II conjugation) microbial metabolites were not impacted, while unconjugated were reduced following Abx, likely representing a reduced enzyme induction resulting from a reduced substrate load.

### Bladder cancer cell proliferation

Urine samples were assayed for their ability to inhibit proliferation in 4 different cancer cell lines (**Figure 3**). Generally, urine appeared to significantly suppress proliferation of bladder cancer cells for all cell lines (observed proliferation values were ≪100% compared to control for these lines) except RT4 (**Figure 3B**). This inhibition of growth could be due to any urine component, including urea, salts, native flavanols and flavanol microbial metabolites (note that urines samples were processed prior to application to reduce urea and salt levels). SCaBER and SW870 (**Figure 3A, C**) cells appeared to be more sensitive to urine overall, and interestingly seemed to have both greater variability (SEM) for all groups and greater separation between control and Abx treatments. The nature of the flavanol did not have a significant impact on proliferation but administration of Abx significantly increased proliferation (in terms of main factor effects; no differences were observed between Control and Abx for specific flavonoids). The lack of an effect of C/EC or GSE compared to vehicle in urines from animals not administered Abx was somewhat surprising, considering the high levels of native flavanols and flavanol microbial metabolites in the C/EC or GSE urines compared to vehicle (**Figure 2A-B**). However, the large inter-animal variability observed in native flavanols, valerolactones, phenylalkyl acids (**Figure 2A, C, D**) and other flavanol microbial metabolites may have contributed to subsequent inter-animal variability in the cell lines that were sensitive to treatment (SCaBER and SW870).

**Figure 3.**
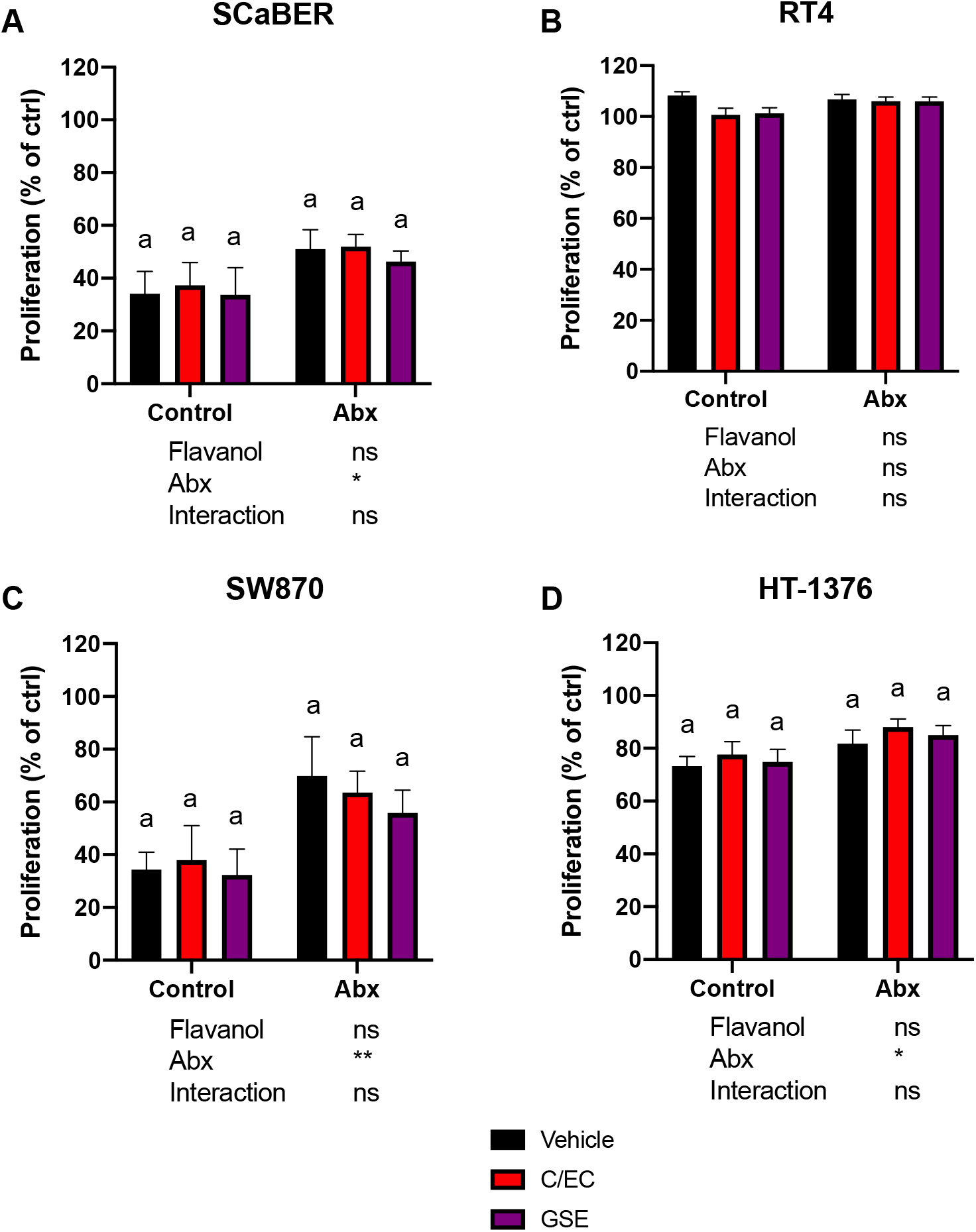
Impact of urine (10% in media) on proliferation in various bladder cancer cell lines (expressed as a % relative to water control). Values are presented as mean ± SEM. For each cell line, 2-way ANOVA was performed to determine the statistical significance of main effects (flavanol and Abx treatment) and interactions. Values below each graph indicate the results of 2-way ANOVA. If a significant main effect or interaction was detected, Holm-Sidak post hoc tests to account for multiple comparisons were performed to determine differences among the 3 flavanol treatments within each antibiotic treatment group (Control and Abx); bars not sharing a common superscript letter within each group are statistically different. Holm-Sidak post hoc tests were also performed to determine differences between antibiotic treatments (Control and Abx) for each flavanol; asterisks indicate a significant difference for that flavanol between Control and Abx (\**P*<0.05, \*\**P*<0.01, \*\*\**P*<0.001, \*\*\*\**P*<0.0001). The overall family-wise error rate was set as 0.05, with one family per group.

It is interesting that administration of Abx appeared to increase proliferation across all treatments. This could be ascribed to the Abx themselves increasing proliferation, or the absence of microbial products that may reduce proliferation. A previous study demonstrated that Abx are cytotoxic to bladder cancer cells(Kamat & Lamm 2004), which is the opposite of what we observed. Therefore, it is worth further investigation to determine the contribution of background microbial metabolites (arising from typical diets, not specifically flavanols) to bladder cancer proliferation. Although not part of the original hypothesis, this finding may suggest that an intact microbiome provides protection against uncontrolled growth of bladder cancer cells.

### Associations between composition and activity

Due to the lack of clear treatment effects from flavanols on proliferation, as well as high inter-animal variability observed in compounds expected to have anti-proliferative activity [native flavanols(Bai & Wang 2012; Hazafa et al. 2020) and their microbial metabolites(Álvarez-Cilleros et al. 2018)], we next performed an individual animal-level analysis independent of treatment groups. We determined which urine samples were the most vs. least effective inhibitors of proliferation (greatest vs. least % proliferation compared to control), irrespective of treatment group, and selected the most effective vs. least effective quartiles (7 animals per quartile). Unsurprisingly, there were clear highly significant separations, with low variability, between the upper and lower quartiles of efficacy in each cell line (**Figure 4A**). Interestingly, the separations between the upper and lower quartiles of efficacy were more pronounced in magnitude for SCaBER and SW870 than for RT4 and HT-1376, similar to what was observed in the treatment group analysis (**Figure 3**). Again, these data suggest that these cells are more sensitive to treatment, and molecular differentiation of these cell lines may provide insight into the mechanisms behind these differences.

**Figure 4.**
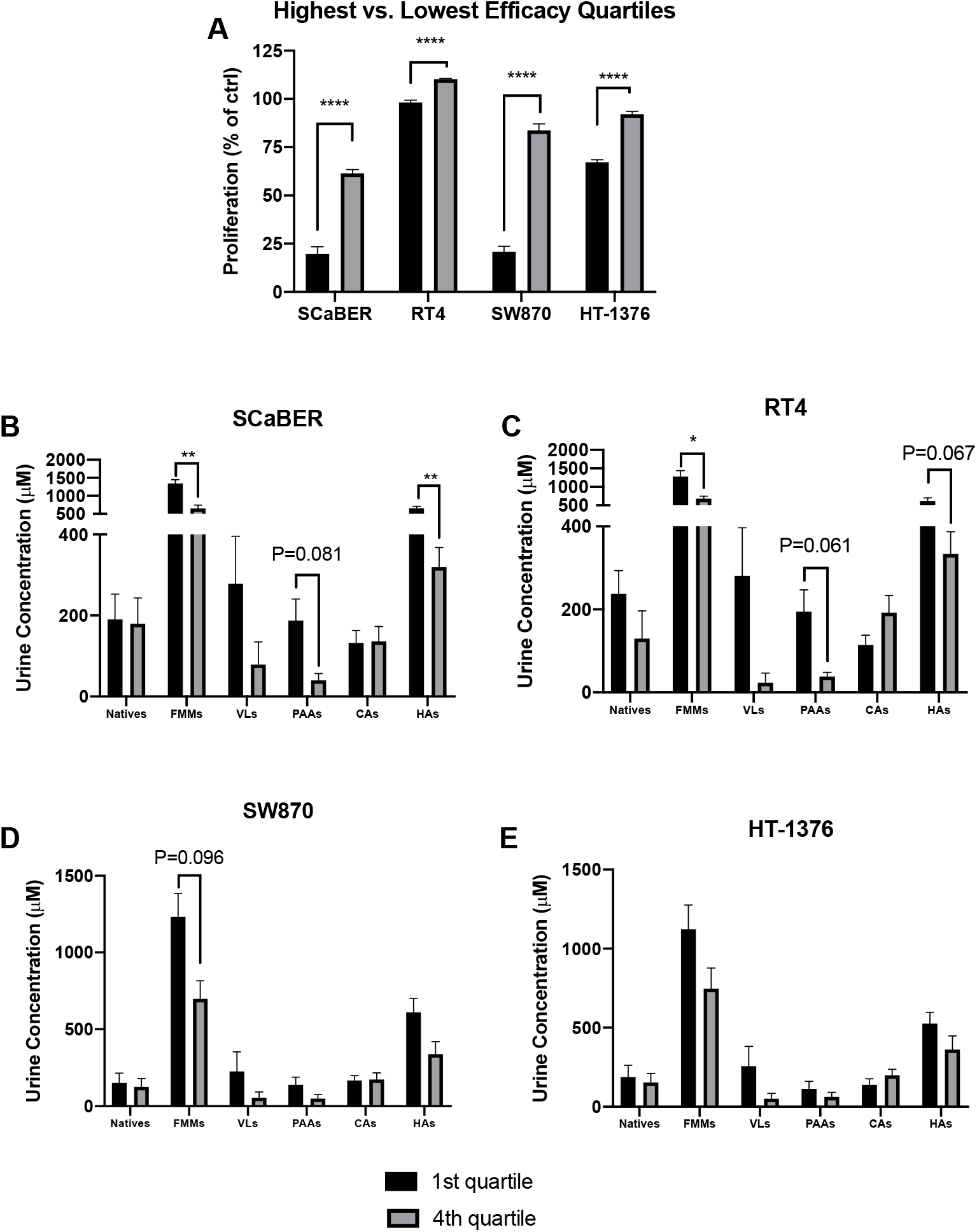
**A**) Differentiation of urine samples by the highest vs. lowest quartiles of anti-proliferative activity (lowest vs. highest % proliferation compared to water control) for various bladder cancer cell lines. Values are presented as mean ± SEM. Proliferation was compared between the highest vs. lowest quartiles by t-tests, with the Holm-Sidak method to control for multiple comparisons (1 family, 4 t-tests: 1 per cell line), with the overall family α= 0.05. Each cell line was analyzed individually, without assuming a consistent SD. **B-E**) Comparison of urine compositions between the most effective vs. least effective samples for each cell line. Values are presented as mean ± SEM. Within each cell line, levels of each compound class were compared between the highest vs. lowest quartiles by t-tests, with the Holm-Sidak method to control for multiple comparisons (1 family per cell line, 6 t-tests: 1 per compound class). For all tests: \**P*<0.05, \*\**P*<0.01, \*\*\**P*<0.001, \*\*\*\**P*<0.0001.

These individual-level differences provided sufficient separation and low enough variability to then attempt to differentiate urine compositional factors associated with enhanced vs. reduced efficacy that were not observable at the treatment levels. We thus compared the urine composition of the most vs. least effective urine samples by compound class and cell line (**Figure 4B-E**). Benzoic acids, other aromatics and non-aromatics were excluded due to lack of significant differences across treatments (**Figure 2G-I**).

In SCaBER cells (**Figure 4B**), significantly higher levels of total flavanol microbial metabolites and hippuric acids (P=0.002 and 0.003, respectively), and borderline significantly higher levels of phenylalkyl acids (P=0.081) were found in the most effective urines compared to the least effective urines. In RT4 cells (**Figure 4C**), significantly higher levels of total flavanol microbial metabolites, and borderline significantly higher levels of phenylalkyl acids (P=0.061) and hippuric acids (0.067) were found in the most effective urines compared to the least effective urines. In SW870 cells (**Figure 4D**), no significant differences were found between the most effective urines compared to the least effective urines, however the differences in phenylalkyl acids were borderline significantly higher (P=0.096). In HT-1376 cells (**Figure 4E**), no significant or borderline significant differences were found between the most effective urines compared to the least effective urines. It warrants mention that due to the exploratory nature of this analysis, these statistics were conservative, in that the overall family (1 family per cell line, 6 t-test per family) was set at α= 0.05 to reduce the rate of false positives; adjusted P-values are presented and used for evaluation of statistical significance. Although not statistically significant due to the conservative statistical approach, generally total flavanol microbial metabolites, valerolactones, phenylalkyl acids and hippuric acids appeared to trend universally toward higher levels in the most effective urine samples. Valerolactones likely would have been statistically different in SCaBER and RT4 cells if not for the large variability in the most effective samples. Interestingly, native flavanols levels (which are typically viewed as the main bioactives delivered to target tissues following consumption by researchers studying flavanol-rich foods), did not appear to be associated with enhanced efficacy in any of the 4 cell lines. This is an intriguing finding, suggesting that flavanol microbial metabolites warrant more consideration as bioactive compounds arising from flavanol compounds. Cinnamic acid levels also did not seem related to efficacy, perhaps unsurprisingly due to the lack of difference among flavanol groups (**Figure 2E**).

The findings in **Figure 4** suggest that certain compound classes are associated with enhanced anti-proliferative activity of bladder cancer cells, when the highest vs. lowest efficacy urine samples were selectively compared. In order to further evaluate these findings, the reverse analysis was also performed: the highest vs. lowest quartile of each compound class were determined, and the anti-proliferative activities of these urine samples were compared (again, benzoic acids, other aromatics and non-aromatics were excluded, **Figure 5**).

**Figure 5.**
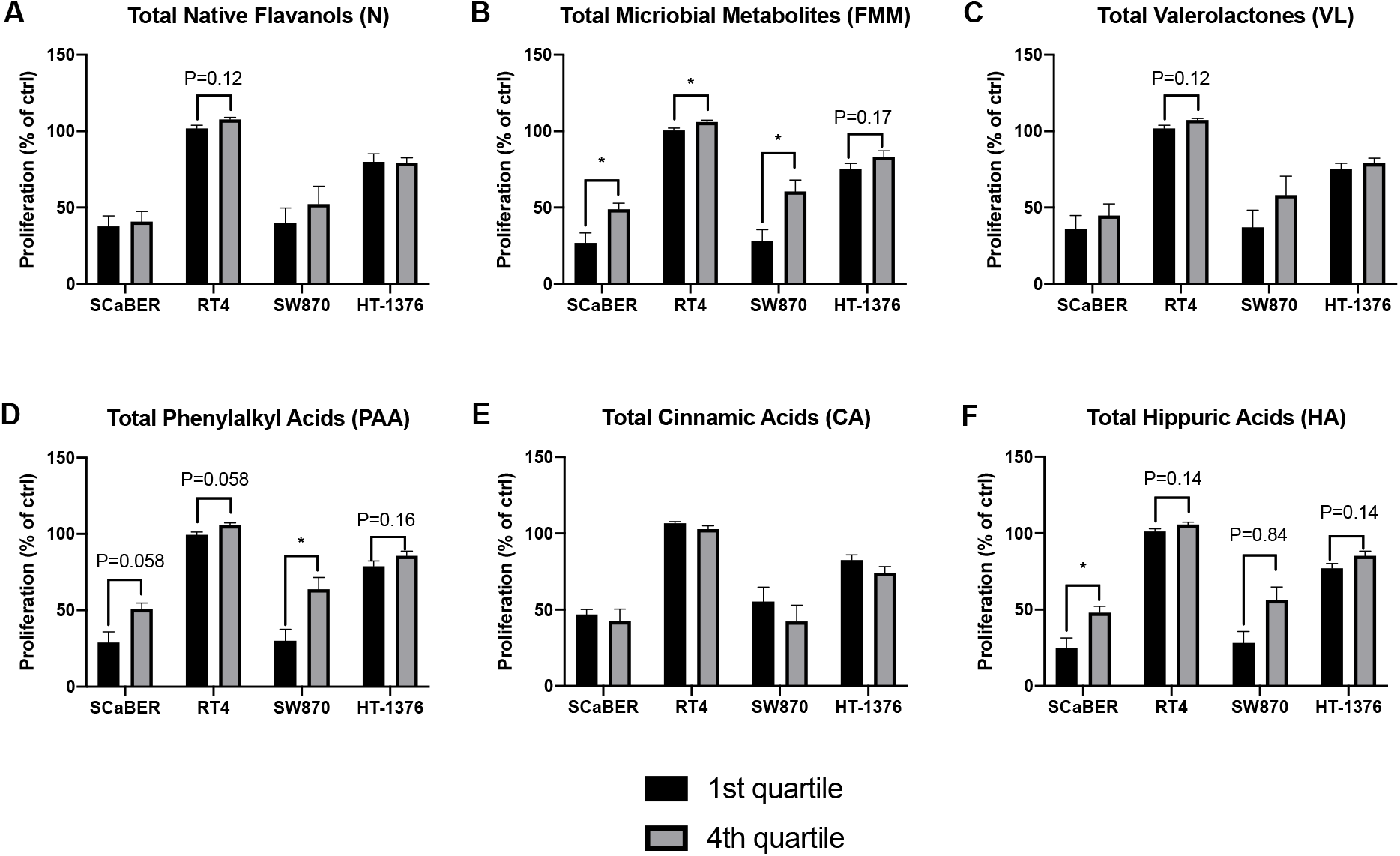
Comparison of proliferation between urine samples with the highest and lowest quartiles of concentrations of various classes of compounds. Values are presented as mean ± SEM. Within each compound class, proliferation was compared between the highest vs. lowest quartiles by t-tests, with the Holm-Sidak method to control for multiple comparisons (1 family per compound class, 4 t-tests: 1 per cell line). For all tests: \**P*<0.05, \*\**P*<0.01, \*\*\**P*<0.001, \*\*\*\**P*<0.0001.

No statistical significance in proliferation between the highest and lowest quartiles of native flavanols was detected in any cell line (**Figure 5A**). The only borderline significant difference observed was for RT4 cells (P=0.12). Significantly lower levels of proliferation were observed in the highest quartiles compared to the lowest for all cell lines except HT-1376 (**Figure 5B**), for which a borderline significance (P=0.17) was detected. No statistical significance in proliferation between the highest and lowest quartiles of valerolactones was detected in any cell line (**Figure 5C**). The only borderline significant difference observed was for RT4 cells (P=0.12). For phenylalkyl acids, a significant difference between the highest and lowest quartiles was observed in SW870 cells (P=0.035, **Figure 5D**), and borderline significant differences were observed for SCaBER, RT4 and HT-1376 cells (P=0.058, 0.058, and 0.16, respectively). For cinnamic acids, no significant or borderline significant differences in proliferation between the highest and lowest quartiles were detected in any cell line (**Figure 5E**). For hippuric acids, a significant difference between the highest and lowest quartiles was observed in SCaBER cells (P=0.039, **Figure 5F**), and borderline significant differences were observed for SW870, RT4 and HT-1376 cells (P=0.14, 0.084, and 0.14, respectively). Again, to reduce false positives, the statistical analysis was conservative: the overall family (1 family compound, 4 t-tests per cell line) was set at α= 0.05. Although not as ideal as validating the findings from **Figure 4** in a separate dataset, the findings presented in Figure 5 provide further evidence that higher levels of total flavanol microbial metabolites, and certain classes of flavanol microbial metabolites (particularly phenylalkyl acids and hippuric acids) are broadly associated with enhanced inhibition of bladder cancer cell proliferation (and vice versa) *in vitro*. Again, SCaBER and SW870 cells appeared to be more sensitive to these compounds (or other factors in the corresponding urine samples) than RT4 and HT-1376 cell lines.

### Characteristics of cell models

To begin to explore why our different cell lines differentially responded to the microbial metabolite-containing urines, we first explored previously published drug sensitivities(Seashore-Ludlow et al. 2015). This data set included SCaBER, RT4, and HT-1376 cells screened with 481 FDA-approved drugs, clinical candidates, and small molecule probes. In general, SCaBER cells are more sensitive than RT4 or HT-1376 cells to drug treatments, (**Supplemental Table 4**), and thus differing responses to other xenobiotics such as flavanols and their metabolites is not surprising. We also examined whether our responder cell lines (SCaBER and SW780) and non-responder cell lines (RT4 and HT-1376) shared mutations, as we reasoned it was plausible that non-responder cells had mutations making them resistant or responder cell lines had mutations that sensitizes these cell lines to microbial metabolites. We first identified mutations in each of the four cell lines using the Cancer Cell Line Encyclopedia(Barretina et al. 2012; Cancer Cell Line Encyclopedia Consortium & Genomics of Drug Sensitivity in Cancer Consortium 2015; Ghandi et al. 2019; Li et al. 2019; The Broad Institute). We identified 20 shared genes that were mutated in SCaBER and SW780 cells, and 11 genes between RT4 and HT-1376 cells (**Supplemental Table 5**). Between these two data sets, only *TTN* was shared. To examine the frequency of mutation in these genes in human bladder cancer patients, we examined 412 muscle invasive bladder cancer cases from The Cancer Genome Atlas(Robertson et al. 2017). Here, we observed deletions and amplification of shared genes at variable frequencies (**Supplemental Figure 1**). Whether these mutations contribute to sensitivity or resistance of our microbial metabolites remains to be experimentally determined, but these observations further reinforce the concept that factors such as patient and tumor genetics, tumor type and stage, etc. could be used to design personalized adjuvant therapy regimens.

### Discussion

To our knowledge, this study is novel in that it is the first to examine the role that the gut microbiome relationships between flavanols, the gut microbiome, and bladder cancer proliferation using physiologically relevant mixtures of flavanol microbial metabolites. *Ex vivo* studies utilizing mouse serum and cardiovascular cells has been employed previously, and were used as a model for our unique approach(Petersen et al. 2018). As mentioned previously, there is evidence that flavanol microbial metabolites (rather than native flavanols) may be primarily responsible for the bioactivities of these compounds in peripheral tissues due to their predominance in circulation, however, studies to date have been limited to *in vitro* models using purified single sources of flavanol microbial metabolites or basic mixtures of chemically synthesized compounds(Fernandez-Millan et al. 2014; Bitner et al. 2018). The present study design allowed for the generation of physiologically relevant mixtures of native flavanols and flavanol microbial metabolites for use in subsequent cell culture models, while also demonstrating the active role of the gut microbiome in the generation of flavanol microbial metabolites with bioactivities potentially more potent than native compounds in some cases. Moreover, the experimental design used here demonstrates the interindividual variability in metabolism and circulation of flavanols/flavanol microbial metabolites from dietary sources, which would not be possible using traditional *in vitro* approaches with purified flavanol compounds. This approach allowed for identification of individual animals as responders and non-responders and did not rely on the assumption that all individuals in a group would respond similarly. Moreover, the interindividual variability in circulating levels of flavanols/flavanol microbial metabolites in a study where all other aspects of diet are controlled support the notion that utilizing phytochemicals in medicinal applications should be a personalized approach. A natural extension of this approach is that the nature of the tumor (as observed based on differences in the responses for each cell line), as well as the individual genetics of the patient (polymorphisms in nuclear receptors, detoxification pathways, etc. that govern flavonoid activity, absorption, and excretion) are likely to play a role in the benefits of flavanols (and generally, flavonoids and other polyphenols) and microbial metabolites for cancer prevention and adjuvant therapy. We have recently begun initial investigations toward understanding the impact of genetics of the effects of flavonoids on disease outcomes(Griffin et al. Submitted).

While differences between the 4 cell lines employed were somewhat expected, the observation that SCaBER and SW870 cells were overall more sensitive to all treatments (**Figure 3**), and displayed greater differences in proliferation between the highest and lowest quartiles in response to treatment (**Figure 4A**), is worth noting. Bladder cancer tumors are typically heterogeneous, with non-muscle invasive tumors generally showing similar molecular and genetic subtypes that differ from muscle invasive tumors(Choi et al. 2014; McConkey & Choi 2018). Within muscle invasive bladder cancer, luminal subtype tumors are predictive of greater overall survival, however, certain basal subtypes are more responsive to neoadjuvant chemotherapy(Choi et al. 2014; Seiler et al. 2017; McConkey & Choi 2018). To account for these two general subtypes in our studies, we have utilized two basal and two luminal bladder cancer cell lines. There is no common origin between these cell lines, as SW870 are luminal cells from a female patient, and SCaBER are basal cells from a male patient as mentioned above. Moreover, SW780 cells have previously been demonstrated to be more sensitive to flavanols (Philips et al. 2009; Luo et al. 2017).

Further investigation into genetic variation and response to flavanols and their metabolites are warranted. Using flavanols and their metabolites in various bladder cancer cell lines, chemically-induced cancer in animals with genetic diversity (collaborative-cross and diversity outbred models), and eventually cohorts of human subjects will reveal the importance of genetics and sensitivity to treatment. From these studies, correlations between genetics and response to treatment can be used to home in on genetic loci, and even specific genes, that dictate sensitivity to flavonoids, by strategies such as QTL mapping and GWAS studies. Such data could then be used to profile potential patients for optimizing treatment strategies.

Although we were able to demonstrate that the gut microbiome plays a role in bladder cancer cell proliferation with this model, it is important to note several limitations. Firstly, we selected urine to capture bioavailable compounds arising from flavanol consumption, as urine samples represent the major excretion pathway for these compounds and collection over 48 hours represents chronic exposure as opposed to a single time point. While utilizing flavanols from urine *in vitro* in peripheral tissue cell culture would be unorthodox in most cases, it does have particular relevance in this case given that we studied uroepithelial cells, which are naturally exposed to urine. The urine was processed after collection to precipitate urea and salts, but it was not possible to achieve complete sample purity. It is possible that residual urea and salt concentrations could have impacted the results of the study; solid-phase extraction could be used in future studies to reduce this potential issue. Secondly, while this method utilized physiologically relevant mixtures of flavanols and their microbial metabolites, it was not possible to identify individual compounds with anti-proliferative activities. Rather, it was possible to identify urine profiles and classes of compounds with enhanced activity – notably valerolactones, phenylalkyl acids and hippuric acids. As bladder cancer occurs more frequently in men than women, we only used male rats. Whether there are differences in microbial metabolite processing between male and female rats is presently unknown, but should be addressed experimentally. Finally, it must be mentioned that GSE is comprised of predominantly dimers and oligomers. GSE was selected due to its purity and history of producing beneficial effects in metabolic studies(Goodrich et al. 2012; Griffin et al. 2017). However, it is possible that utilizing a flavanol source with a larger concentration of polymeric and thus poorly bioavailable flavanols (such as cocoa extract) would have generated greater differences in the urine concentrations of total flavanols, native flavanols, and flavanol microbial metabolites in the presence and absence of Abx, which could ultimately yield greater differences in cell culture assays. This possibility will be explored in future studies.

Given that the results of this study suggest that flavanol microbial metabolites may exert anti-proliferative effects in bladder cancer cells, future efforts should be directed at utilizing this model in other cell lines to further assess flavanol microbial metabolite bioactivity compared to native flavanols. Further efforts to identify specific flavanol microbial metabolites related to anti-proliferation of bladder cancer cells should also be performed, so that specific flavanols may ultimately be used as an adjuvant to traditional bladder cancer therapies. Additionally, it would be beneficial to study the anti-bladder cancer effect of flavanols utilizing germ-free rodent models to control for microbiome and Abx effects.

## CONCLUSIONS

Urine represents a major accumulation site for dietary flavanols and their microbial and Phase-II metabolites prior to elimination. Bladder epithelial cells are therefore exposed to high concentrations of these compounds for long periods of time, suggesting the possibility for significant protective effects in the bladder arising from flavanol consumption. Microbial metabolites of flavanols (particularly valerolactones, phenylalkyl acids and hippuric acids) are associated with reduced bladder cancer cell proliferation *in vitro*. Significant interindividual variability in urine profiles resulting from flavanol intake were observed within treatment groups, despite the use of inbred rats fed a common background diet. Furthermore, the responses of various bladder cancer cell lines differed, along with significant interindividual variability in the anti-proliferative activities of urine compounds within treatment groups. These results highlight the potential for enhancing our understanding of individual genetics, microbiome profiles, flavonoid metabolism profiles, tumor characteristics, etc. in order to design personalized dietary interventions for cancer prevention and/or adjuvant therapy to reduce bladder cancer incidence and improve outcomes. Such possibilities justify the need for future studies using an individualized approach to research, where animals or subjects are investigated on an individual basis in addition to grouped aggregates.

## ACKNOLWEDGEMENT

Funding for this project was provided through startup funding from North Carolina State University, as well as support from the North Carolina Agricultural Research Service (NCARS) and the Hatch Program of the National Institute of Food and Agriculture, U.S. Department of Agriculture (APN). We would also like to acknowledge the Case Research Institute, a joint venture between University Hospitals and Case Western Reserve University, start-up funds (to MMG), and the Cell and Molecular Biology Training Program (T32 GM 008056 to SEK).

## Notes

### Competing Interest Statement

The authors have declared no competing interest.

