## Supplementary Information for "Microbial metabolites of flavanols in urine are associated with enhanced anti-proliferative activity in bladder cancer cells in vitro"

Supplementary Information
for

***Microbial metabolites of flavanols in urine are associated with enhanced anti-proliferative activity in bladder cancer cells in vitro***

Laura E. Griffin[a], Sarah E. Kohrt[b,c], Atul Rathore[a], Colin D. Kay[a], Magdalena M. Grabowska[b,c,d,e], Andrew P. Neilson[a]*

[a]Plants for Human Health Institute, Department of Food, Bioprocessing and Nutrition Sciences, North Carolina State University, Kannapolis, NC
[b]Case Comprehensive Cancer Center, Case Western Reserve University, Cleveland, OH
[c]Department of Pharmacology, Case Western Reserve University, Cleveland, OH
[d]Department of Urology, Case Western Reserve University, Cleveland, OH
[e]Department of Biochemistry, Case Western Reserve University, Cleveland, OH

*Corresponding author: Dr. Andrew Neilson,, Plants for Human Health Institute, Department of Food, Bioprocessing and Nutrition Sciences, North Carolina State University, 600 Laureate Way, Kannapolis, NC 28081, 704-250-5495

| **Supplementary Table 1.** Reference standard list and number of qualitative MS transitions utilized[1] | | | | | | |
|---|---|---|---|---|---|---|
| **Full Name[1]** | **Abbreviation** | **CAS** | **Reference Standard** | **Relative Quantifier** | **MRM Transitions** | **Ionization mode** |
| 3,4-dihydroxyphenyl glycol | 34-diOH-PGOL | 28822-73-3 | ✓ | reference standard | 5 | − |
| hydroxy(4-hydroxy-3-methoxyphenyl)acetic acid | 4-OH-3-OCH-MndA | 55-10-7 | ✓ | reference standard | 5 | − |
| 1-(4-hydroxy-3-methoxyphenyl)-1,2-ethanediol | 4-OH-3-OCH-PheEGlyc | 534-82-7 | ✓ | reference standard | 5 | − |
| 3-hydroxybenzoic acid-4-O-glucuronide | 3-OH-BA-4-OGlcA | synthetic (UEA) | ✓ | reference standard | 4 | − |
| 2-amino-3-(4-hydroxy-3-methoxyphenyl)propanoic acid | 2-NH2-4-OH-PPA | 7636-26-2 | ✓ | reference standard | 5 | − |
| (3,4-dihydroxyphenyl)-2-hydroxyacetaldehyde | DOPEGAL | 13023-73-9 | ✓ | reference standard | 5 | − |
| 5-hydroxybenzoic acid-3-sulfate | 5-OH-BA-3-Sulf | synthetic (TRC[2] D451710) | ✓ | reference standard | 2 | − |
| 3-hydroxyphenethyl-4-sulfate | 3-OH-PEtOH-4-Sulf | synthetic (TRC H949705) | ✓ | reference standard | 5 | − |
| 4-hydroxyphenylethanol-3-sulfate | 4-OH-PEtOH-3-sulf | 1391053-88-5 | ✓ | reference standard | 5 | − |
| 4-hydroxybenzoic acid-3-O-glucuronide | 4-OH-BA-3-OGlcA | synthetic (UEA) | ✓ | reference standard | 3 | − |
| 5-hydroxyphenylpropanoic acid-3-O-glucuronide | 5-OH-PPA-3-OGlcA | synthetic (TRC D454370) | ✓ | reference standard | 3 | − |
| hydroxyhippuric acid_Isomer1 | OH-HA_IS1 | x | x | 4-hydroxyhippuric acid | 3 | − |
| 4-hydroxyhippuric acid | 4-OH-HA | 2482-25-9 | ✓ | reference standard | 3 | − |
| (3-hydroxyphenyl)-3-hydroxypropanoic acid | 3-OH-3-OH-PPA | 3247-75-4 | ✓ | reference standard | 5 | − |
| 3-hydroxyhippuric acid | 3-OH-HA | 1637-75-8 | ✓ | reference standard | 4 | − |
| hydroxyphenyl-gamma-valerolactone-O-glucuronide | OH-phenyl-g-Val-OGlcA | x | x | 3-hydroxyphenyl-gamma-valerolactone-4-sulfate | 6 | − |
| 4-methoxybenzoic acid-3-O-glucuronide | 4-OCH-BA-3-OGlcA | synthetic (UEA) | ✓ | reference standard | 3 | − |
| 4-hydroxyphenylpropanoic acid-3-sulfate | 4-OH-PPA-3-Sulf | 1187945-70-5 | ✓ | reference standard | 4 | − |
| 4-hydroxybenzoic acid-4-sulfate | 4-OH-BA-Sulf | synthetic (UEA) | ✓ | reference standard | 2 | − |
| 3,5-dihydroxybenzaldehyde | 35-diOH-Bald | 26153-38-8 | ✓ | reference standard | 2 | − |
| hippuric acid | HA | 495-69-2 | ✓ | reference standard | 3 | − |
| 3-methoxyphenylacetic acid-4-sulfate | 3-OCH-PAA-4-Sulf | 38339-06-9 | ✓ | reference standard | 5 | − |
| 5-(phenyl)-gamma-valerolactone-3-sulfate-4-O-glucuronide | phenyl-Val-Sulf-OGlcA | x | x | 3-hydroxyphenyl-gamma-valerolactone-4-sulfate | 5 | − |
| 3-hydroxybenzaldehyde | 3-OH-Bald | 100-83-4 | ✓ | reference standard | 3 | − |
| 3-methoxyphenylpropanoic acid-4-sulfate | 3-OCH-PPA-4-Sulf | 86321-33-7 | ✓ | reference standard | 5 | − |
| hydroxybenzenediol-sulfate | OH-Bz-Sulf | x | x | 5-hydroxybenzoic acid-3-sulfate | 4 | − |
| (−)-N-benzoylglutamic acid | N-benzyglut A | 6094-36-6 | ✓ | reference standard | 4 | − |
| procyanidin B1 | Procyanidin B1 | 20315-25-7 | ✓ | reference standard | 5 | − |
| hydroxyphenylvalerolactone-sulfate | OH-phenyl-Val-Sulf | x | x | 3-hydroxyphenyl-gamma- | 4 | − |

| | | | | | | |
|---|---|---|---|---|---|---|
| | | | | valerolactone-4-sulfate | | |
| epicatechin sulf-O-glucuronide_Isomer1 | Ecat-Sulf-OGlcA_IS1 | x | x | (−)-epicatechin-3′-sulfate | 3 | − |
| 1-methoxybenzene-2-O-glucuronide | 1-OCH-Bz-2-OGlcA | 111897-99-5 | ✓ | reference standard | 3 | − |
| 5-(4’-hydroxyphenyl)-gamma-valerolactone-3’-sulfate | 4-OH-phenyl-g-Val-3-Sulf | synthetic in-house | ✓ | reference standard | 4 | − |
| 3-hydroxyphenyl-gamma-valerolactone-4-sulfate | 3-OH-phenyl-g-Val-4-Sulf | synthetic (TRC D454570) | ✓ | reference standard | 5 | − |
| 5-(4'-hydroxyphenyl)-gamma-valerolactone-3'-O-glucuronide | 4-OH-phenyl-g-Val-3-OGlcA | synthetic in-house | ✓ | reference standard | 1 | − |
| 5-(3'-hydroxyphenyl)-gamma-valerolactone-4'-O-glucuronide | 3-OH-phenyl-g-Val-4-OGlcA | synthetic in-house | ✓ | reference standard | 2 | − |
| 3-hydroxy-4-methoxyphenylacetic acid | 3-OH-4-OCH-PAA | 1131-94-8 | ✓ | reference standard | 2 | − |
| 5-(4',5'-dihydroxyphenyl)-gamma-valerolactone-O-sulfate_Isomer2 | diOH-phenyl-Val-Sulf_IS2 | x | x | 3-hydroxyphenyl-gamma-valerolactone-4-sulfate | 4 | − |
| 3,4-dihydroxyphenyl-gamma-valerolactone | 34-diOH-phenyl-g-Val | 191666-22-5 | ✓ | reference standard | 6 | − |
| methoxycinnamic acid-O-glucuronide | OCH-CIA-OGlcA | x | x | 4-methoxycinnamic acid-3-O-glucuronide | 4 | − |
| 5-(4',5'-dihydroxyphenyl)-gamma-valerolactone-O-sulfate_Isomer1 | diOH-phenyl-Val-Sulf_IS1 | x | x | 3-hydroxyphenyl-gamma-valerolactone-4-sulfate | 4 | − |
| 4-methoxycinnamic acid-3-O-glucuronide | 4-OCH-CIA-3-OGlcA | 1065272-10-7 | ✓ | reference standard | 6 | − |
| epicatechin sulf-O-glucuronide_Isomer2 | Ecat-Sulf-OGlcA_IS2 | x | x | (−)-epicatechin-3′-sulfate | 3 | − |
| 4-methoxycinnamic acid-3-sulfate | 4-OCH-CIA-3-Sulf | synthetic (TRC I816160) | ✓ | reference standard | 3 | − |
| 4-hydroxycinnamic acid | 4-OH-CIA | 7400-08-0 | ✓ | reference standard | 3 | − |
| cinnamic acid-4-sulfate | CIA-4-Sulf | 308338-96-7 | ✓ | reference standard | 4 | − |
| 4-hydroxy-3-methoxybenzaldehyde | 4-OH-3-OCH-Bald | 121-33-5 | ✓ | reference standard | 5 | − |
| 3-hydroxyphenylpropanoic acid | 3-OH-PPA | 621-54-5 | ✓ | reference standard | 2 | − |
| epicatechin sulf-O-glucuronide_Isomer3 | Ecat-Sulf-OGlcA_IS3 | x | x | (−)-epicatechin-3′-sulfate | 3 | − |
| 3-hydroxy-4-methoxybenzaldehyde | 3-OH-4-OCH-Bald | 621-59-0 | ✓ | reference standard | 3 | − |
| 4-hydroxyphenylpropanoic acid | 4-OH-PPA | 501-97-3 | ✓ | reference standard | 3 | − |
| procyanidin B2 | Procyanidin B2 | 29106-49-8 | ✓ | reference standard | 5 | − |
| 4-methylhippuric acid | 4-M-HA | 27115-50-0 | ✓ | reference standard | 5 | − |
| hydroxyhippuric acid_Isomer2 | OH-HA_IS2 | x | x | 4-hydroxyhippuric acid | 3 | − |
| (−)-epicatechin-3′-sulfate | Ecat-3-Sulf | 1038922-77-8 | ✓ | reference standard | 6 | − |
| (−)-epicatechin-3′-O-glucuronide | Ecat-3-OGlcA | synthetic in-house | ✓ | reference standard | 5 | − |
| 4-hydroxy-3-methoxyphenylpropanoic acid | 4-OH-3-OCH-PPA | 1135-23-5 | ✓ | reference standard | 5 | − |
| 3-hydroxy-4-methoxyphenylpropanoic acid | 3-OH-4-OCH-PPA | 1135-15-5 | ✓ | reference standard | 3 | − |

| | | | | | | |
|---|---|---|---|---|---|---|
| 5-(3',4'-dihydroxyphenyl)-gamma-valerolactone | diOH-phenyl-Val | x | x | (−)-epicatechin-3′-O-glucuronide | 5 | − |
| (−)-epicatechin | Ecat | 490-46-0 | ✓ | reference standard | 6 | − |
| (3-hydroxy-4-methoxyphenyl)ethanone | 3-OH-4-OCH_Phe_ethanone | 6100-74-9 | ✓ | reference standard | 5 | − |
| 3'-O-methyl-(-)-epicatechin-5-sulfate | 3M-Ecat-5-Sulf | synthetic in-house | ✓ | reference standard | 6 | − |
| 3'-O-methyl-(-)-epicatechin-4-sulfate | 3M-Ecat-4-Sulf | synthetic in-house | ✓ | reference standard | 8 | − |
| 5-(4'-hydroxyphenyl)-gamma-valerolactone | 4-OH-phenyl-Val | 871329-30-5 | ✓ | reference standard | 3 | − |
| 4-hydroxy-3,5-dimethoxybenzaldehyde | 4-OH-35-diOCH-Bald | 134-96-3 | ✓ | reference standard | 5 | − |
| 3'-O-methyl-(-)-epicatechin-7-sulfate | 3M-Ecat-7-Sulf | synthetic in-house | ✓ | reference standard | 6 | − |
| 4'-O-methyl-(-)-epicatechin-5-sulfate | 4M-Ecat-5-Sulf | synthetic in-house | ✓ | reference standard | 7 | − |
| 5-(3'-hydroxyphenyl)-gamma-valerolactone | 3-OH-phenyl-Val | 21618-91-7 | ✓ | reference standard | 4 | − |
| procyanidin C1 | Procyanidin C1 | 37064-30-5 | ✓ | reference standard | 6 | − |
| 4′-O-methyl(−)-epicatechin-7-sulfate | 4M-Ecat-7-Sulf | synthetic in-house | ✓ | reference standard | 7 | − |
| 4-hydroxy-3-methoxycinnamaldehyde | 4-OH-3-OCH-CIAld | 458-36-6 | ✓ | reference standard | 5 | − |
| 5-(4-hydroxyphenyl)valeric acid | 4-OH-phenyl-ValA | 4654-08-4 | ✓ | reference standard | 5 | − |
| 5-phenylvaleric acid | 5-Phen-ValA | 2270-20-4 | ✓ | reference standard | 5 | − |
| 4-hydroxy-3,5-dimethoxybenzoic acid | 4-OH-35-diOCH-BA | 530-57-4 | ✓ | reference standard | 3 | + |
| 4-hydroxy-3,5-dimethoxyphenylacetic acid | 4-OH-35-diOCH-PAA | 4385-56-2 | ✓ | reference standard | 5 | + |
| 3,4,5-trihydroxybenzoic acid-sulfate | 345-triOH-BA-Sulf | x | x | 5-hydroxybenzoic acid-3-sulfate | 5 | + |

[1]Ordered by retention time within ionization mode (+ or -)
[2]Toronto Research Chemicals (TRC; 20 Martin Ross Avenue, North York, ON Canada, M3J 2K8; Website: https://www.trc-canada.com/)

| **Supplementary Table 2.** Classification of compounds detected in rat urine samples | | | |
|---|---|---|---|
| **Native or metabolite** | **Metabolite class** | **Conjugation state** | **Compound name** |
| native | N/A | unconj. | (epi)catechin |
| | | conj. | (−)-epicatechin-3′-O-glucuronide |
| | | conj. | (−)-epicatechin-3′-sulfate |
| | | conj. | 3'-O-methyl-(-)-epicatechin-5-sulfate/3'-O-methyl-(-)-epicatechin-4'-sulfate |
| | | conj. | 3'-O-methyl-(-)-epicatechin-7-sulfate |
| | | conj. | 4'-O-methyl-(-)-epicatechin-5-sulfate |
| | | conj. | 4′-O-methyl(−)-epicatechin-7-sulfate |
| | | unconj. | procyanidin B1 |
| | | unconj. | procyanidin B2 |
| | | unconj. | procyanidin C1 |
| | | conj. | epicatechin sulfoglucuronide isomer |
| | | conj. | epicatechin sulfoglucuronide isomer |
| | | conj. | epicatechin sulfoglucuronide isomer |
| | | unconj. | procyanidin B2 |
| microbial metabolite | benzoic acid deriv. | unconj. | 4-hydroxy-3,5-dimethoxybenzaldehyde |
| | | unconj. | 4-hydroxy-3-methoxybenzaldehyde |
| | | conj. | 3-hydroxybenzoic acid-4-O-glucuronide |
| | | conj. | 4-methoxybenzoic acid-3-O-glucuronide |
| | | conj. | 4-hydroxybenzoic acid-4-sulfate |
| | | conj. | 1-methoxybenzene-2-O-glucuronide |
| | | unconj. | 3,5-dihydroxybenzaldehyde |
| | | unconj. | 3-hydroxy-4-methoxybenzaldehyde |
| | | conj. | 4-hydroxybenzoic acid-3-O-glucuronide |
| | | unconj. | (-)-N-benzoylglutamic Acid |
| | | unconj. | 3-hydroxybenzaldehyde |
| | | unconj. | 4-hydroxy-3,5-dimethoxybenzoic acid |
| | | conj. | 3,4,5-trihydroxy benzoic acid sulfate |
| | cinnamic acid deriv. | unconj. | 4-hydroxy-3-methoxycinnamaldehyde |
| | | unconj. | 4-hydroxycinnamic acid |
| | | conj. | cinnamic acid-4-sulfate |
| | | conj. | 4-methoxycinnamic acid-3-O-Glucuronide |
| | | conj. | 4-methoxycinnamic acid-3-sulfate |
| | | conj. | methoxycinnamic acid-O-glucuronide |
| | hippuric acid deriv. | unconj. | 4-hydroxyhippuric acid |
| | | unconj. | 3-hydroxyhippuric acid |
| | | unconj. | 4-methylhippuric acid |
| | | unconj. | hippuric acid |
| | | unconj. | hydroxyhippuric acid_Isomer3 |
| | | unconj. | hydroxyhippuric acid_Isomer1 |
| | non-aromatic | unconj. | oxalic acid |
| | other aromatics | unconj. | 3,4-dihydroxyphenyl glycol |
| | | unconj. | (3-Hydroxy-4-methoxyphenyl)ethanone |
| | | unconj. | 1-(4-hydroxy-3-methoxyphenyl)-1,2-ethanediol |
| | | conj. | 3-hydroxyphenethyl-4-sulfate |
| | | conj. | 4-hydroxyphenylethanol-3-sulfate |
| | | conj. | hydroxybenzenediol-sulfate isomer |
| | phenylalkyl acid deriv. | unconj. | 5-(4-hydroxyphenyl)valeric acid |
| | | unconj. | (3-hydroxyphenyl)-3-hydroxypropanoic acid |
| | | unconj. | 2-amino-3-(4-hydroxy-3-methoxyphenyl)propanoic acid |
| | | unconj. | (3,4-dihydroxyphenyl)-2-hydroxyacetaldehyde |
| | | unconj. | 4-hydroxy-3-methoxyphenyl(hydroxy)acetic acid |
| | | unconj. | 5-phenylvaleric acid |
| | | unconj. | 4-hydroxyphenylpropanoic acid |
| | | conj. | 5-hydroxyphenylpropanoic acid-3-O-glucuronide |
| | | unconj. | 4-hydroxy-3-methoxyphenylpropanoic acid/3-hydroxy-4-methoxyphenylacetic acid |
| | | conj. | 3-methoxyphenylacetic acid-4-sulfate |

| | | | |
|---|---|---|---|
| | | unconj. | 3-hydroxy-4-methoxyphenylacetic acid |
| | | unconj. | 3-hydroxyphenylpropanoic acid |
| | | conj. | 4-hydroxyphenylpropanoic acid-3-Sulfate |
| | | conj. | 5-hydroxybenzoic acid-3-sulfate |
| | | conj. | 3-methoxyphenylpropanoic acid-4-sulfate |
| | | unconj. | 4-hydroxy-3,5-dimethoxyphenylacetic acid |
| | valerolactone deriv. | unconj. | 3,4-dihydroxyphenyl-gamma-valerolactone |
| | | unconj. | 5-(4'-hydroxyphenyl)-gamma-valerolactone |
| | | unconj. | 5-(3'-hydroxyphenyl)-gamma-valerolactone |
| | | conj. | 3-hydroxyphenyl-gamma-valerolactone-4-sulfate |
| | | conj. | 5-(4’-hydroxyphenyl)-γ –valerolactone-3’-sulfate |
| | | conj. | 5-(3'-hydroxyphenyl)-gamma-valerolactone-4'-O-glucuronide |
| | | conj. | 5-(4'-hydroxyphenyl)-γ-valerolactone-3'-O-glucuronide |
| | | conj. | 5-(Phenyl)-γ-valerolactone-3-sulfate-4-O-glucuronide |
| | | conj. | 5-(4',5'-dihydroxyphenyl)-gamma-valerolactone-O-sulfate Isomer 1 |
| | | conj. | 5-(4',5'-dihydroxyphenyl)-gamma-valerolactone-O-sulfate Isomer 2 |
| | | unconj. | 5-(3',4'-dihydroxyphenyl)-gamma-valerolactone |
| | | conj. | hydroxyphenyl-gamma-valerolactone-O-glucuronide |
| | | conj. | hydroxyphenylvalerolactone-sulfate |

**Supplementary Table 3.** Concentrations ($\mu$M) of analytes in individual volume-normalized urine samples. Compounds detected but below the linear range were reported as 0.0001 $\mu$M. Compounds not detected were reported as 0 $\mu$M

| | **Treatment (Flavanol and Abx)** | | | | | | | | | | | | | | | | | | | | | | | | | | | |
|---|---|---|---|---|---|---|---|---|---|---|---|---|---|---|---|---|---|---|---|---|---|---|---|---|---|---|---|---|
| | **C/EC** | **C/EC** | **C/EC** | **C/EC** | **C/EC** | **C/EC** | **C/EC** | **C/EC** | **C/EC** | **C/EC** | **GSE** | **GSE** | **GSE** | **GSE** | **GSE** | **GSE** | **GSE** | **GSE** | **GSE** | **GSE** | **Vehicle** | **Vehicle** | **Vehicle** | **Vehicle** | **Vehicle** | **Vehicle** | **Vehicle** | **Vehicle** |
| **Compound** | + | + | + | + | + | − | − | − | − | − | + | + | + | + | + | − | − | − | − | − | + | + | + | + | − | − | − | − |
| (epi)catechin | 116.97 | 78.25 | 20.74 | 109.71 | 78.21 | 35.91 | 97.75 | 27.62 | 206.26 | 105.06 | 41.71 | 85.37 | 100.31 | 22.77 | 6.99 | 143.78 | 241.25 | 245.07 | 17.19 | 101.45 | 0.45 | 0.52 | 0.49 | 0.59 | 0.53 | 0.69 | 0.56 | 0.45 |
| (−)-epicatechin-3′-O-glucuronide | 5.89 | 13.68 | 2.29 | 15.33 | 14.24 | 2.47 | 3.95 | 4.50 | 12.50 | 6.75 | 4.44 | 4.46 | 11.13 | 3.25 | 2.37 | 12.41 | 20.99 | 16.02 | 0.67 | 4.80 | 0.01 | 0.00 | 0.00 | 0.00 | 0.02 | 0.02 | 0.01 | 0.01 |
| (−)-epicatechin-3′-sulfate | 5.16 | 6.40 | 0.21 | 12.53 | 11.43 | 0.48 | 0.21 | 0.55 | 4.89 | 3.69 | 0.15 | 0.51 | 9.54 | 0.36 | 0.12 | 8.35 | 2.41 | 4.29 | 0.16 | 0.64 | 0.00 | 0.00 | 0.00 | 0.00 | 0.00 | 0.00 | 0.01 | 0.00 |
| 3'-O-methyl-(-)-epicatechin-5-sulfate/3'-O-methyl-(-)-epicatechin-4'-sulfate | 15.24 | 0.00 | 0.00 | 29.36 | 28.37 | 0.00 | 0.00 | 3.43 | 12.42 | 9.08 | 0.00 | 0.00 | 40.42 | 0.00 | 0.00 | 19.18 | 8.86 | 11.90 | 1.23 | 3.21 | 0.01 | 0.00 | 0.01 | 0.00 | 0.00 | 0.00 | 0.02 | 0.02 |
| 3'-O-methyl-(-)-epicatechin-7-sulfate | 57.68 | 77.92 | 11.28 | 83.35 | 101.35 | 11.63 | 15.19 | 14.44 | 63.12 | 40.06 | 17.19 | 14.96 | 130.26 | 13.88 | 12.88 | 47.26 | 63.21 | 59.84 | 3.66 | 14.64 | 0.05 | 0.03 | 0.03 | 0.03 | 0.03 | 0.04 | 0.04 | 0.02 |
| 4'-O-methyl-(-)-epicatechin-5-sulfate | 63.07 | 87.40 | 12.10 | 97.79 | 93.10 | 13.02 | 16.48 | 15.88 | 68.97 | 39.12 | 18.29 | 17.09 | 145.29 | 15.64 | 13.94 | 47.46 | 67.21 | 63.27 | 4.03 | 17.19 | 0.07 | 0.04 | 0.04 | 0.03 | 0.05 | 0.08 | 0.05 | 0.02 |
| 4′-O-methyl(−)-epicatechin-7-sulfate | 0.01 | 0.19 | 0.01 | 0.14 | 0.23 | 0.00 | 0.00 | 0.00 | 0.11 | 0.10 | 0.01 | 0.02 | 0.21 | 0.01 | 0.01 | 0.00 | 0.00 | 0.14 | 0.01 | 0.01 | 0.00 | 0.00 | 0.00 | 0.00 | 0.00 | 0.00 | 0.00 | 0.00 |
| procyanidin B1 | 0.02 | 0.02 | 0.04 | 0.03 | 0.03 | 0.08 | 0.04 | 0.06 | 0.02 | 0.01 | 0.25 | 0.20 | 0.19 | 0.10 | 0.03 | 0.04 | 0.03 | 0.03 | 0.02 | 0.03 | 0.02 | 0.02 | 0.02 | 0.02 | 0.01 | 0.01 | 0.02 | 0.01 |
| procyanidin B2 | 0.03 | 0.03 | 0.10 | 0.03 | 0.03 | 0.16 | 0.08 | 0.18 | 0.03 | 0.03 | 0.72 | 0.40 | 0.48 | 0.18 | 0.06 | 0.03 | 0.03 | 0.03 | 0.03 | 0.16 | 0.03 | 0.03 | 0.03 | 0.03 | 0.03 | 0.03 | 0.03 | 0.03 |
| procyanidin C1 | 0.04 | 0.04 | 0.04 | 0.04 | 0.04 | 0.04 | 0.04 | 0.04 | 0.04 | 0.04 | 0.04 | 0.04 | 0.00 | 0.00 | 0.04 | 0.04 | 0.00 | 0.04 | 0.04 | 0.04 | 0.04 | 0.04 | 0.04 | 0.00 | 0.04 | 0.04 | 0.04 | 0.04 |
| epicatechin sulfoglucuronide isomer 1 | 0.05 | 0.61 | 0.10 | 0.40 | 0.58 | 0.13 | 0.20 | 0.28 | 0.47 | 0.14 | 0.06 | 0.01 | 0.38 | 0.09 | 0.11 | 0.65 | 0.64 | 0.74 | 0.06 | 0.19 | 0.11 | 0.06 | 0.07 | 0.10 | 0.05 | 0.09 | 0.07 | 0.02 |
| epicatechin sulfoglucuronide isomer 2 | 0.00 | 0.00 | 0.00 | 0.00 | 0.00 | 0.00 | 0.00 | 0.00 | 0.00 | 0.00 | 0.00 | 0.00 | 0.00 | 0.13 | 0.09 | 0.00 | 0.00 | 0.00 | 0.00 | 0.12 | 0.00 | 0.01 | 0.02 | 0.00 | 0.01 | 0.00 | 0.01 | 0.00 |
| epicatechin sulfoglucuronide isomer 3 | 0.00 | 0.00 | 0.00 | 0.00 | 0.00 | 0.00 | 0.00 | 0.00 | 0.00 | 0.00 | 0.00 | 0.00 | 0.00 | 0.12 | 0.10 | 0.00 | 0.00 | 0.00 | 0.00 | 0.00 | 0.00 | 0.01 | 0.01 | 0.00 | 0.01 | 0.00 | 0.02 | 0.00 |
| procyanidin B2 | 0.04 | 0.04 | 0.13 | 0.04 | 0.04 | 0.16 | 0.10 | 0.19 | 0.05 | 0.04 | 0.92 | 0.49 | 0.55 | 0.29 | 0.09 | 0.04 | 0.04 | 0.06 | 0.05 | 0.12 | 0.04 | 0.04 | 0.04 | 0.04 | 0.04 | 0.04 | 0.04 | 0.04 |
| 4-hydroxy-3,5-dimethoxybenzaldehyde | 0.04 | 0.03 | 0.03 | 0.03 | 0.03 | 0.04 | 0.04 | 0.07 | 0.03 | 0.03 | 0.03 | 0.02 | 0.02 | 0.04 | 0.07 | 0.02 | 0.07 | 0.03 | 0.02 | 0.03 | 0.03 | 0.04 | 0.02 | 0.04 | 0.03 | 0.04 | 0.03 | 0.03 |
| 4-hydroxy-3-methoxybenzaldehyde | 0.00 | 0.09 | 0.11 | 0.10 | 0.04 | 0.04 | 0.11 | 0.05 | 0.32 | 0.14 | 0.15 | 0.09 | 0.04 | 0.0001 | 0.05 | 0.08 | 0.10 | 0.16 | 0.06 | 0.14 | 0.12 | 0.04 | 0.02 | 0.0001 | 0.07 | 0.04 | 0.06 | 0.09 |
| 3-hydroxybenzoic acid-4-O-glucuronide | 0.04 | 0.11 | 0.14 | 0.09 | 0.08 | 0.15 | 0.14 | 0.24 | 0.03 | 0.03 | 0.13 | 0.13 | 0.08 | 0.18 | 0.14 | 0.03 | 0.07 | 0.03 | 0.10 | 0.17 | 0.14 | 0.05 | 0.08 | 0.07 | 0.10 | 0.07 | 0.09 | 0.12 |
| 4-methoxybenzoic acid-3-O-glucuronide | 0.12 | 0.45 | 0.86 | 0.16 | 0.21 | 0.69 | 0.89 | 0.82 | 0.21 | 0.15 | 0.85 | 0.57 | 0.31 | 0.71 | 0.73 | 0.09 | 0.09 | 0.09 | 1.09 | 0.40 | 0.34 | 0.11 | 0.23 | 0.21 | 0.55 | 0.27 | 0.19 | 0.64 |
| 4-hydroxybenzoic acid-4-sulfate | 57.88 | 65.70 | 66.21 | 53.78 | 55.03 | 58.38 | 83.83 | 62.01 | 55.99 | 38.81 | 53.85 | 76.39 | 69.56 | 48.47 | 44.40 | 34.00 | 52.61 | 55.47 | 44.18 | 99.61 | 47.10 | 97.64 | 51.94 | 60.19 | 52.02 | 59.64 | 107.23 | 83.97 |
| 1-methoxybenzene-2-O-glucuronide | 0.05 | 0.06 | 0.05 | 0.00 | 0.04 | 0.00 | 0.00 | 0.00 | 0.00 | 0.00 | 0.07 | 0.04 | 0.04 | 0.04 | 0.04 | 0.00 | 0.00 | 0.00 | 0.00 | 0.00 | 0.03 | 0.04 | 0.00 | 0.08 | 0.00 | 0.00 | 0.10 | 0.00 |
| 3,5-dihydroxybenzaldehyde | 0.10 | 0.20 | 0.30 | 0.13 | 0.22 | 0.20 | 0.22 | 0.17 | 0.34 | 0.31 | 0.11 | 0.05 | 0.62 | 0.04 | 0.12 | 1.21 | 0.53 | 0.39 | 0.20 | 0.46 | 0.07 | 0.45 | 0.14 | 0.12 | 0.21 | 0.25 | 0.25 | 0.27 |
| 3-hydroxy-4-methoxybenzaldehyde | 0.01 | 0.15 | 0.07 | 0.04 | 0.04 | 0.18 | 0.26 | 0.16 | 0.28 | 0.17 | 0.00 | 0.00 | 0.00 | 0.03 | 0.03 | 0.11 | 0.27 | 0.21 | 0.04 | 0.11 | 0.08 | 0.03 | 0.03 | 0.04 | 0.04 | 0.03 | 0.06 | 0.00 |
| 4-hydroxybenzoic acid-3-O-glucuronide | 0.02 | 0.02 | 0.09 | 0.02 | 0.03 | 0.12 | 0.16 | 0.10 | 0.06 | 0.02 | 0.12 | 0.09 | 0.03 | 0.09 | 0.08 | 0.04 | 0.06 | 0.05 | 0.11 | 0.15 | 0.03 | 0.02 | 0.02 | 0.03 | 0.09 | 0.04 | 0.05 | 0.07 |
| (-)-N-benzoylglutamic Acid | 0.55 | 0.80 | 1.10 | 0.65 | 0.70 | 0.77 | 1.09 | 0.92 | 0.49 | 0.33 | 0.57 | 0.78 | 0.97 | 0.59 | 0.63 | 0.44 | 0.92 | 0.63 | 0.91 | 0.94 | 0.48 | 0.92 | 0.57 | 0.74 | 1.06 | 0.62 | 0.94 | 1.03 |
| 3-hydroxybenzaldehyde | 2.60 | 1.67 | 1.82 | 1.06 | 1.18 | 1.84 | 2.58 | 1.64 | 2.58 | 1.06 | 2.15 | 2.38 | 1.21 | 1.44 | 1.54 | 1.08 | 1.81 | 1.65 | 1.93 | 3.76 | 2.56 | 2.88 | 1.46 | 2.19 | 3.13 | 2.34 | 1.81 | 2.95 |
| 4-hydroxy-3,5-dimethoxybenzoic acid | 0.19 | 0.58 | 0.64 | 0.19 | 0.25 | 0.16 | 0.32 | 0.37 | 0.16 | 0.08 | 0.65 | 0.37 | 0.33 | 0.47 | 0.55 | 0.06 | 0.09 | 0.10 | 0.08 | 0.33 | 0.44 | 0.33 | 0.28 | 0.26 | 0.13 | 0.11 | 0.14 | 0.10 |
| 4-hydroxy-3-methoxycinnamaldehyde | 0.01 | 0.0001 | 0.0001 | 0.01 | 0.00 | 0.0001 | 0.01 | 0.0001 | 0.01 | 0.00 | 0.00 | 0.0001 | 0.00 | 0.0001 | 0.0001 | 0.00 | 0.02 | 0.01 | 0.0001 | 0.00 | 0.0001 | 0.00 | 0.0001 | 0.0001 | 0.00 | 0.0001 | 0.0001 | 0.00 |
| 4-hydroxycinnamic acid | 63.95 | 163.71 | 231.73 | 65.92 | 62.12 | 67.07 | 73.47 | 70.19 | 18.09 | 7.61 | 134.68 | 120.23 | 169.47 | 113.30 | 144.65 | 1.60 | 7.10 | 4.31 | 30.06 | 79.57 | 187.29 | 152.47 | 101.74 | 106.11 | 68.28 | 39.19 | 97.69 | 105.09 |
| cinnamic acid-4-sulfate | 63.00 | 182.26 | 251.86 | 71.74 | 70.00 | 95.58 | 100.21 | 99.79 | 23.77 | 17.05 | 144.17 | 132.90 | 205.17 | 124.43 | 142.63 | 21.01 | 61.88 | 37.24 | 46.52 | 99.42 | 210.00 | 165.06 | 107.00 | 127.20 | 77.38 | 43.65 | 107.37 | 110.00 |
| 4-methoxycinnamic acid-3-O-Glucuronide | 0.85 | 4.98 | 3.93 | 1.16 | 1.89 | 1.01 | 1.21 | 1.34 | 0.27 | 0.55 | 2.66 | 1.72 | 2.45 | 1.36 | 2.58 | 0.87 | 0.80 | 0.36 | 0.68 | 1.01 | 3.18 | 0.61 | 2.15 | 1.22 | 1.46 | 0.57 | 0.94 | 0.90 |
| 4-methoxycinnamic acid-3-sulfate | 0.07 | 0.30 | 0.36 | 0.08 | 0.12 | 0.24 | 0.44 | 0.53 | 1.20 | 0.88 | 0.16 | 0.19 | 0.25 | 0.10 | 0.23 | 2.08 | 2.51 | 2.79 | 0.38 | 0.65 | 0.29 | 0.17 | 0.11 | 0.09 | 0.09 | 0.05 | 0.14 | 0.10 |
| methoxycinnamic acid-O-glucuronide Isomer 3 | 0.52 | 2.88 | 2.48 | 0.72 | 1.04 | 0.51 | 0.88 | 1.08 | 0.56 | 0.62 | 1.42 | 1.26 | 1.51 | 0.75 | 1.73 | 1.34 | 1.98 | 1.67 | 0.52 | 0.78 | 1.86 | 0.42 | 1.34 | 0.95 | 0.83 | 0.35 | 0.69 | 0.65 |
| 4-hydroxyhippuric acid | 11.16 | 12.00 | 15.85 | 16.19 | 11.62 | 14.02 | 15.02 | 11.92 | 10.05 | 9.68 | 13.44 | 13.17 | 14.93 | 22.88 | 15.03 | 8.47 | 6.52 | 7.74 | 14.07 | 13.70 | 17.13 | 16.47 | 11.42 | 12.52 | 13.09 | 9.53 | 16.96 | 19.09 |
| 3-hydroxyhippuric acid | 0.21 | 0.41 | 0.47 | 0.23 | 0.23 | 46.69 | 68.88 | 55.38 | 17.14 | 11.01 | 0.27 | 0.35 | 0.40 | 0.19 | 0.38 | 8.00 | 49.91 | 34.76 | 56.67 | 32.21 | 0.34 | 0.46 | 0.43 | 0.30 | 1.97 | 1.70 | 3.21 | 2.79 |
| 4-methylhippuric acid | 0.68 | 0.91 | 0.58 | 0.78 | 0.82 | 0.55 | 0.58 | 0.67 | 0.37 | 0.64 | 0.74 | 0.85 | 0.55 | 0.53 | 0.63 | 0.58 | 0.35 | 0.25 | 0.46 | 0.59 | 0.48 | 0.57 | 0.36 | 0.64 | 0.75 | 0.59 | 0.80 | 0.73 |
| hippuric acid | 166.86 | 199.98 | 231.48 | 205.17 | 199.80 | 488.61 | 691.61 | 777.91 | 371.81 | 368.93 | 169.56 | 244.71 | 255.30 | 214.22 | 208.04 | 630.74 | 713.29 | 493.94 | 704.35 | 672.36 | 183.34 | 243.41 | 169.00 | 234.46 | 444.40 | 356.29 | 610.15 | 511.42 |
| hydroxyhippuric acid_Isomer3 | 4.52 | 7.53 | 14.40 | 5.84 | 4.95 | 15.43 | 26.16 | 22.69 | 5.93 | 4.80 | 16.62 | 14.50 | 6.58 | 10.66 | 10.09 | 6.33 | 10.68 | 8.71 | 17.40 | 20.14 | 7.50 | 8.01 | 5.13 | 6.74 | 11.77 | 7.43 | 15.94 | 10.44 |
| hydroxyhippuric acid_Isomer1 | 2.52 | 2.60 | 3.02 | 3.50 | 2.49 | 3.15 | 3.60 | 2.79 | 2.41 | 2.13 | 3.35 | 3.09 | 3.15 | 4.82 | 3.42 | 2.03 | 1.61 | 2.10 | 3.38 | 3.37 | 3.65 | 3.47 | 2.46 | 2.94 | 3.11 | 2.37 | 3.79 | 4.32 |
| oxalic acid | 11.87 | 12.64 | 11.41 | 12.83 | 12.54 | 13.47 | 12.18 | 11.89 | 13.19 | 12.61 | 11.66 | 12.72 | 13.31 | 12.42 | 13.36 | 12.29 | 11.40 | 12.94 | 12.13 | 11.57 | 12.16 | 13.15 | 12.23 | 12.79 | 13.06 | 12.83 | 11.99 | 13.18 |
| 3,4-dihydroxyphenyl glycol | 0.02 | 0.03 | 0.01 | 0.03 | 0.03 | 0.05 | 0.04 | 0.02 | 0.0001 | 0.06 | 0.07 | 0.03 | 0.18 | 0.04 | 0.0001 | 0.06 | 0.04 | 0.01 | 0.01 | 0.09 | 0.07 | 0.12 | 0.01 | 0.09 | 0.06 | 0.04 | 0.18 | 0.06 |
| (3-Hydroxy-4-methoxyphenyl)ethanone | 30.08 | 0.00 | 0.71 | 0.00 | 0.00 | 2.02 | 2.53 | 3.55 | 0.00 | 0.00 | 0.16 | 0.00 | 91.36 | 0.29 | 0.63 | 0.00 | 0.00 | 0.00 | 2.39 | 3.36 | 1.20 | 1.07 | 0.55 | 0.73 | 0.61 | 0.92 | 1.64 | 1.48 |
| 1-(4-hydroxy-3-methoxyphenyl)-1,2-ethanediol | 0.08 | 0.07 | 0.00 | 0.05 | 0.07 | 0.10 | 0.14 | 0.08 | 0.00 | 0.01 | 0.00 | 0.15 | 0.04 | 0.12 | 0.18 | 0.06 | 0.00 | 0.00 | 0.02 | 0.00 | 0.01 | 0.13 | 0.03 | 0.11 | 0.14 | 0.10 | 0.15 | 0.14 |
| 3-hydroxyphenethyl-4-sulfate | 0.04 | 0.03 | 0.07 | 0.06 | 0.06 | 0.09 | 0.05 | 0.03 | 0.03 | 0.02 | 0.03 | 0.09 | 0.05 | 0.18 | 0.12 | 0.03 | 0.02 | 0.01 | 0.22 | 0.09 | 0.06 | 0.11 | 0.06 | 0.07 | 0.13 | 0.08 | 0.07 | 0.19 |
| 4-hydroxyphenylethanol-3-sulfate | 0.04 | 0.03 | 0.07 | 0.06 | 0.06 | 0.09 | 0.05 | 0.03 | 0.03 | 0.02 | 0.03 | 0.08 | 0.05 | 0.18 | 0.12 | 0.03 | 0.02 | 0.01 | 0.23 | 0.09 | 0.06 | 0.11 | 0.06 | 0.07 | 0.13 | 0.08 | 0.08 | 0.19 |
| urolithin A | 0.00 | 0.00 | 0.0001 | 0.00 | 0.00 | 0.02 | 0.05 | 0.01 | 0.03 | 0.01 | 0.00 | 0.00 | 0.00 | 0.00 | 0.0001 | 0.01 | 0.03 | 0.04 | 0.02 | 0.06 | 0.00 | 0.01 | 0.0001 | 0.00 | 0.01 | 0.04 | 0.05 | 0.04 |
| hydroxybenzenediol-sulfate isomer 2 | 0.03 | 0.03 | 0.03 | 0.02 | 0.03 | 0.06 | 0.04 | 0.06 | 0.03 | 0.01 | 0.03 | 0.04 | 0.03 | 0.02 | 0.03 | 0.04 | 0.03 | 0.02 | 0.03 | 0.05 | 0.02 | 0.04 | 0.03 | 0.03 | 0.06 | 0.04 | 0.04 | 0.07 |
| 5-(4-hydroxyphenyl)valeric acid | 0.0001 | 0.0001 | 0.0001 | 0.0001 | 0.00 | 0.28 | 0.0001 | 0.48 | 0.0001 | 0.0001 | 0.00 | 0.00 | 0.00 | 0.0001 | 0.00 | 0.0001 | 0.07 | 0.0001 | 0.03 | 0.0001 | 0.0001 | 0.0001 | 0.00 | 0.00 | 0.00 | 0.0001 | 0.00 | 0.0001 |
| (3-hydroxyphenyl)-3-hydroxypropanoic acid | 0.0001 | 0.0001 | 0.0001 | 0.0001 | 0.0001 | 0.24 | 0.33 | 0.15 | 0.03 | 0.02 | 0.0001 | 0.0001 | 0.0001 | 0.0001 | 0.0001 | 0.03 | 0.26 | 0.09 | 0.25 | 0.18 | 0.0001 | 0.0001 | 0.0001 | 0.0001 | 0.0001 | 0.0001 | 0.0001 | 0.0001 |
| 2-amino-3-(4-hydroxy-3-methoxyphenyl)propanoic acid | 0.0001 | 0.00 | 0.05 | 0.01 | 0.0001 | 0.02 | 0.0001 | 0.0001 | 0.06 | 0.00 | 0.01 | 0.0001 | 0.00 | 0.03 | 0.02 | 0.00 | 0.01 | 0.0001 | 0.02 | 0.03 | 0.06 | 0.0001 | 0.00 | 0.01 | 0.0001 | 0.0001 | 0.10 | 0.04 |

| | | | | | | | | | | | | | | | | | | | | | | | | | | | | |
|---|---|---|---|---|---|---|---|---|---|---|---|---|---|---|---|---|---|---|---|---|---|---|---|---|---|---|---|---|
| (3,4-dihydroxyphenyl)(hydroxy)acetaldehyde | 12.75 | 30.12 | 10.95 | 12.15 | 14.25 | 16.96 | 8.40 | 4.66 | 5.12 | 7.96 | 0.0001 | 8.00 | 0.0001 | 0.0001 | 0.0001 | 1.73 | 0.0001 | 7.47 | 15.33 | 0.53 | 30.10 | 13.14 | 33.82 | 12.79 | 31.69 | 12.89 | 17.60 | 27.84 |
| 4-hydroxy-3-methoxyphenyl(hydroxy)acetic acid | 0.12 | 0.11 | 0.11 | 0.16 | 0.16 | 0.15 | 0.12 | 0.11 | 0.09 | 0.06 | 0.10 | 0.16 | 0.20 | 0.23 | 0.22 | 0.10 | 0.11 | 0.08 | 0.12 | 0.21 | 0.19 | 0.15 | 0.11 | 0.14 | 0.24 | 0.15 | 0.12 | 0.32 |
| 5-phenyl valeric acid | 0.0001 | 0.06 | 0.0001 | 0.0001 | 0.0001 | 0.10 | 0.94 | 0.06 | 0.0001 | 0.03 | 0.36 | 0.0001 | 0.0001 | 0.13 | 0.0001 | 0.0001 | 0.24 | 0.14 | 0.0001 | 0.0001 | 0.0001 | 0.0001 | 0.0001 | 0.0001 | 0.49 | 0.0001 | 0.0001 | 0.27 |
| 4-hydroxyphenylpropanoic acid | 0.33 | 3.04 | 4.80 | 1.27 | 0.09 | 70.96 | 133.79 | 77.24 | 28.91 | 41.26 | 2.43 | 2.59 | 4.94 | 2.01 | 2.52 | 202.74 | 222.48 | 63.00 | 96.60 | 76.43 | 3.86 | 3.28 | 1.84 | 2.21 | 11.43 | 3.68 | 27.32 | 8.05 |
| 5-hydroxyphenylpropanoic acid-3-O-glucuronide | 0.0001 | 0.01 | 0.04 | 0.00 | 0.0001 | 0.0001 | 0.03 | 0.0001 | 0.03 | 0.03 | 0.0001 | 0.01 | 0.12 | 0.0001 | 0.0001 | 0.25 | 0.12 | 0.02 | 0.0001 | 0.10 | 0.0001 | 0.04 | 0.02 | 0.0001 | 0.01 | 0.0001 | 0.07 | 0.06 |
| 4-hydroxy-3-methoxyphenylpropanoic acid/3-hydroxy-4-methoxyphenylacetic acid | 0.77 | 1.68 | 1.82 | 0.83 | 1.25 | 2.86 | 5.53 | 7.32 | 6.55 | 2.86 | 1.44 | 1.14 | 1.22 | 0.97 | 1.21 | 4.87 | 5.34 | 4.64 | 3.39 | 6.56 | 1.34 | 1.59 | 1.08 | 1.34 | 2.12 | 1.90 | 5.43 | 2.14 |
| 3-methoxyphenylacetic acid-4-sulfate | 0.00 | 0.00 | 0.00 | 0.0001 | 0.00 | 1.46 | 4.57 | 1.34 | 0.30 | 0.13 | 0.11 | 0.00 | 0.00 | 0.00 | 0.00 | 0.57 | 0.54 | 0.26 | 2.00 | 1.78 | 0.00 | 0.00 | 0.00 | 0.00 | 0.37 | 0.17 | 0.98 | 0.47 |
| 3-hydroxy-4-methoxyphenylacetic acid | 0.14 | 1.78 | 1.20 | 0.00 | 0.00 | 0.00 | 0.00 | 0.00 | 0.00 | 0.00 | 0.07 | 0.38 | 0.69 | 0.00 | 0.56 | 0.00 | 0.00 | 0.00 | 0.00 | 0.11 | 1.19 | 0.14 | 0.47 | 0.00 | 0.04 | 0.00 | 0.00 | 0.00 |
| 3-hydroxyphenylpropanoic acid | 0.41 | 0.0001 | 1.59 | 0.0001 | 0.10 | 53.74 | 94.64 | 56.15 | 20.91 | 31.04 | 0.81 | 0.68 | 1.49 | 0.0001 | 0.0001 | 142.65 | 144.95 | 43.17 | 71.07 | 52.08 | 1.28 | 0.89 | 0.62 | 0.0001 | 7.79 | 2.54 | 19.80 | 5.55 |
| 4-hydroxyphenylpropanoic acid-3-Sulfate | 0.00 | 0.09 | 0.00 | 0.00 | 0.00 | 1.46 | 4.52 | 1.31 | 0.30 | 0.15 | 0.14 | 0.00 | 0.00 | 0.00 | 0.00 | 0.58 | 0.50 | 0.29 | 1.93 | 1.73 | 0.00 | 0.00 | 0.00 | 0.00 | 0.40 | 0.17 | 1.00 | 0.43 |
| 5-hydroxybenzoic acid-3-sulfate | 0.06 | 0.04 | 0.54 | 0.06 | 0.07 | 0.24 | 0.16 | 0.14 | 0.03 | 0.02 | 0.29 | 0.54 | 0.08 | 0.75 | 0.49 | 0.03 | 0.01 | 0.01 | 0.24 | 0.23 | 0.06 | 0.16 | 0.09 | 0.09 | 0.13 | 0.08 | 0.09 | 0.14 |
| 3-methoxyphenylpropanoic acid-4-sulfate | 0.39 | 0.73 | 1.00 | 0.44 | 0.41 | 1.25 | 1.68 | 1.38 | 0.79 | 0.37 | 0.73 | 0.76 | 0.66 | 0.70 | 0.73 | 0.73 | 1.26 | 0.90 | 0.79 | 1.66 | 0.70 | 0.70 | 0.59 | 0.57 | 1.41 | 0.41 | 2.96 | 1.09 |
| 4-hydroxy-3,5-dimethoxyphenylacetic acid | 0.11 | 0.34 | 0.24 | 0.08 | 0.14 | 0.04 | 0.05 | 0.08 | 0.08 | 0.04 | 0.16 | 0.16 | 0.20 | 0.11 | 0.17 | 0.04 | 0.05 | 0.07 | 0.04 | 0.05 | 0.29 | 0.11 | 0.13 | 0.12 | 0.05 | 0.04 | 0.20 | 0.06 |
| 3,4-dihydroxyphenyl-gamma-valerolactone | 0.00 | 0.15 | 0.00 | 0.00 | 0.10 | 17.63 | 80.62 | 3.80 | 444.70 | 41.80 | 0.00 | 0.00 | 0.00 | 0.00 | 0.00 | 37.60 | 62.98 | 81.74 | 66.68 | 40.00 | 0.00 | 0.00 | 0.00 | 0.00 | 0.10 | 0.00 | 0.00 | 0.00 |
| 5-(4'-hydroxyphenyl)-gamma-valerolactone | 0.12 | 1.02 | 0.12 | 0.54 | 1.22 | 0.07 | 0.33 | 0.16 | 0.80 | 0.13 | 0.26 | 0.15 | 1.12 | 0.20 | 0.11 | 0.11 | 0.17 | 1.27 | 0.12 | 0.10 | 0.28 | 0.24 | 0.12 | 0.16 | 0.12 | 0.19 | 0.14 | 0.17 |
| 5-(3'-hydroxyphenyl)-gamma-valerolactone | 0.00 | 0.01 | 0.11 | 0.03 | 0.00 | 4.65 | 7.64 | 14.41 | 134.17 | 92.35 | 0.0001 | 0.00 | 0.00 | 0.01 | 0.01 | 114.55 | 110.82 | 159.75 | 19.97 | 21.20 | 0.0001 | 0.0001 | 0.00 | 0.00 | 0.07 | 0.31 | 0.05 | 0.05 |
| 3-hydroxyphenyl-gamma-valerolactone-4-sulfate | 0.08 | 0.19 | 0.27 | 0.09 | 0.11 | 34.97 | 64.70 | 9.97 | 114.97 | 8.99 | 0.05 | 0.14 | 0.14 | 0.05 | 0.17 | 37.84 | 58.44 | 48.28 | 48.15 | 30.47 | 0.17 | 0.07 | 0.19 | 0.07 | 0.07 | 0.11 | 0.07 | 0.05 |
| 5-(4’-hydroxyphenyl)-γ –valerolactone-3’-sulfate | 0.07 | 0.18 | 0.24 | 0.08 | 0.10 | 31.28 | 58.86 | 9.10 | 102.82 | 7.98 | 0.04 | 0.13 | 0.12 | 0.04 | 0.15 | 33.76 | 52.47 | 44.27 | 43.20 | 26.96 | 0.15 | 0.07 | 0.17 | 0.06 | 0.06 | 0.09 | 0.06 | 0.05 |
| 5-(3'-hydroxyphenyl)-gamma-valerolactone-4'-O-glucuronide | 0.0001 | 0.0001 | 0.00 | 0.0001 | 0.0001 | 1.61 | 3.65 | 1.02 | 4.29 | 0.79 | 0.0001 | 0.0001 | 0.0001 | 0.0001 | 0.0001 | 2.63 | 2.95 | 3.85 | 2.79 | 2.76 | 0.0001 | 0.0001 | 0.0001 | 0.0001 | 0.00 | 0.00 | 0.00 | 0.00 |
| 5-(4'-hydroxyphenyl)-γ-valerolactone-3'-O-glucuronide | 0.0001 | 0.0001 | 0.0001 | 0.0001 | 0.0001 | 0.00 | 0.00 | 0.0001 | 4.19 | 0.0001 | 0.0001 | 0.0001 | 0.0001 | 0.0001 | 0.0001 | 0.00 | 0.00 | 0.00 | 0.00 | 0.00 | 0.00 | 0.0001 | 0.0001 | 0.00 | 0.00 | 0.00 | 0.00 | 0.00 |
| 5-(Phenyl)-γ-valerolactone-3-sulfate-4-O-glucuronide | 0.07 | 0.32 | 0.05 | 0.35 | 0.35 | 0.03 | 0.05 | 0.05 | 0.25 | 0.12 | 0.05 | 0.05 | 0.53 | 0.05 | 0.03 | 0.16 | 0.44 | 0.35 | 0.01 | 0.05 | 0.0001 | 0.00 | 0.0001 | 0.00 | 0.00 | 0.0001 | 0.00 | 0.0001 |
| 5-(4',5'-dihydroxyphenyl)-gamma-valerolactone-O-sulfate | 0.00 | 0.01 | 0.03 | 0.00 | 0.01 | 0.00 | 0.00 | 0.00 | 0.00 | 0.00 | 0.02 | 0.02 | 0.03 | 0.02 | 0.01 | 0.00 | 0.08 | 0.00 | 0.00 | 0.00 | 0.02 | 0.02 | 0.01 | 0.01 | 0.01 | 0.00 | 0.01 | 0.02 |
| 5-(4',5'-dihydroxyphenyl)-gamma-valerolactone-O-sulfate | 0.01 | 0.01 | 0.03 | 0.00 | 0.00 | 0.00 | 0.00 | 0.00 | 0.00 | 0.00 | 0.02 | 0.02 | 0.02 | 0.02 | 0.01 | 0.00 | 0.00 | 0.00 | 0.00 | 0.00 | 0.02 | 0.02 | 0.01 | 0.01 | 0.01 | 0.00 | 0.01 | 0.01 |
| 5-(3',4'-dihydroxyphenyl)-gamma-valerolactone | 0.03 | 0.06 | 0.03 | 0.05 | 0.06 | 0.07 | 0.10 | 0.06 | 0.51 | 0.10 | 0.07 | 0.04 | 0.05 | 0.05 | 0.03 | 0.11 | 0.29 | 0.19 | 0.06 | 0.08 | 0.04 | 0.06 | 0.04 | 0.01 | 0.09 | 0.02 | 0.05 | 0.05 |
| hydroxyphenyl-gamma-valerolactone-O-glucuronide_Isomer1 | 0.01 | 0.03 | 0.02 | 0.02 | 0.02 | 0.04 | 0.04 | 0.03 | 0.02 | 0.01 | 0.02 | 0.03 | 0.03 | 0.02 | 0.02 | 0.02 | 0.02 | 0.02 | 0.02 | 0.04 | 0.02 | 0.02 | 0.01 | 0.03 | 0.04 | 0.02 | 0.05 | 0.06 |
| hydroxyphenylvalerolactone-sulfate_Isomer1 | 0.07 | 0.18 | 0.24 | 0.08 | 0.09 | 32.83 | 59.42 | 9.69 | 107.09 | 8.50 | 0.05 | 0.13 | 0.12 | 0.05 | 0.16 | 35.40 | 54.96 | 46.84 | 44.20 | 27.64 | 0.17 | 0.07 | 0.19 | 0.07 | 0.06 | 0.11 | 0.05 | 0.04 |

| **Supplementary Table 4**. Bladder cancer cell line sensitivity to drugs[1] | | | | | |
|---|---|---|---|---|---|
| **Sensitivity[2]** | | | **Compound** | **Gene name of protein target** | **Target or activity of compound** |
| **SCaBER** | **RT4** | **HT-1376** | | | |
| 1.9382 | 9.7431 | 10.082 | oligomycin A | ATP5L2 | inhibitor of mitochondrial ATP synthase |
| 2.3604 | 4.7459 | 10.866 | SB-743921 | KIF11 | inhibitor of kinesin 11 |
| 2.923 | 9.3376 | 12.175 | paclitaxel | | inhibitor of microtubule assembly |
| 3.0442 | 5.1868 | 6.64 | leptomycin B | XPO1 | inhibitor of exportin 1 |
| 3.5435 | 8.3154 | 10.904 | BI-2536 | PLK1 | inhibitor of polo-like kinase 1 (PLK1) |
| 3.5617 | | | dinaciclib | CDK1;CDK2;CDK5;CDK9 | inhibitor of cyclin-dependent kinases |
| 4.7827 | 7.1199 | 6.7813 | LBH-589 | HDAC1;HDAC2;HDAC3;HDAC6;HDAC8 | inhibitor of HDAC1, HDAC2, HDAC3, HDAC6, and HDAC8 |
| 5.08 | 5.4913 | 7.217 | ouabain | ATP1A1;ATP1A2;ATP1A3;ATP1A4;ATP1B1;ATP1B2;ATP1B3;ATP1B4 | cardiac glycoside; inhibitor of the Na+/K+-ATPase |
| 5.3538 | 11.074 | 11.879 | BRD-K97651142 | | aphrocallistin derivative |
| 5.6955 | 7.2359 | 10.816 | GSK461364 | PLK1 | inhibitor of polo-like kinase 1 (PLK1) |
| 5.8447 | 7.5297 | 11.225 | SNX-2112 | HSP90AA1;HSP90B1 | inhibitor of HSP90alpha and HSP90beta |
| 5.8898 | 9.0936 | 10.362 | afatinib | EGFR;ERBB2 | inhibitor of EGFR and HER2 |
| 5.9274 | | | AT13387 | HSP90AA1 | inhibitor of HSP90 |
| 6.2984 | 9.6557 | 11.848 | KX2-391 | SRC | peptide mimetic; inhibitor of SRC activity in cells |
| 6.3105 | 9.1778 | 10.217 | 1S,3R-RSL-3 | GPX4 | synthetic lethal with HRAS in engineered cells; inhibitor of GPX4 |
| 6.381 | 9.3434 | 12.073 | rigosertib | PIK3CA;PIK3CB;PLK1 | inhibitor of polo-like kinase 1; inhibitor of PI3K catalytic subunits alpha and beta |
| 6.4544 | 8.4468 | 10.961 | vincristine | | inhibitor of mictrotubule assembly |
| 6.5894 | 10.441 | 8.8831 | dasatinib | EPHA2;KIT;LCK;SRC;YES1 | inhibitor of SRC, YES1, EPHA2, c-KIT, and LCK |
| 6.7611 | 6.8773 | 10.028 | omacetaxine mepesuccinate | | inhibitor of protein translation by preventing protein elongation |
| 6.919 | | | alvocidib | CDK1;CDK2;CDK4;CDK6 | inhibitor of cyclin-dependent kinases |
| | 4.6395 | 9.3541 | CR-1-31B | EIF4A2;EIF4E;EIF4G1 | silvestrol analog; inhibits translation by modulating the eIF4F complex |
| 2.3604 | 4.7459 | 10.866 | SB-743921 | KIF11 | inhibitor of kinesin 11 |
| 3.0442 | 5.1868 | 6.64 | leptomycin B | XPO1 | inhibitor of exportin 1 |
| 10.109 | 5.2954 | 10.916 | brefeldin A | ARF1 | modulator of ADP-ribosylation factor 1; inhibitor of protein translocation from ER to Golgi |
| 5.08 | 5.4913 | 7.217 | ouabain | ATP1A1;ATP1A2;ATP1A3;ATP1A4;ATP1B1;ATP1B2;ATP1B3;ATP1B4 | cardiac glycoside; inhibitor of the Na+/K+-ATPase |
| 6.7611 | 6.8773 | 10.028 | omacetaxine mepesuccinate | | inhibitor of protein translation by preventing protein elongation |
| 4.7827 | 7.1199 | 6.7813 | LBH-589 | HDAC1;HDAC2;HDAC3;HDAC6;HDAC8 | inhibitor of HDAC1, HDAC2, HDAC3, HDAC6, and HDAC8 |
| 7.1592 | 7.2261 | 10.617 | doxorubicin | TOP2A | inhibitor of topoisomerase II |
| 5.6955 | 7.2359 | 10.816 | GSK461364 | PLK1 | inhibitor of polo-like kinase 1 (PLK1) |
| 7.7222 | 7.3893 | 9.9091 | SR-II-138A | EIF4A2;EIF4E;EIF4G1 | silvestrol analog; inhibits translation by modulating the eIF4F complex |
| 5.8447 | 7.5297 | 11.225 | SNX-2112 | HSP90AA1;HSP90B1 | inhibitor of HSP90alpha and HSP90beta |
| 11.93 | 8.0839 | 13.914 | clofarabine | | inducer of DNA damage |
| 3.5435 | 8.3154 | 10.904 | BI-2536 | PLK1 | inhibitor of polo-like kinase 1 (PLK1) |
| 7.4058 | 8.3521 | 9.7719 | MLN2238 | PSMB5 | inhibitor of 20S proteasome at the chymotrypsin-like proteolytic (beta-5) site |
| 8.8724 | 8.3558 | 12.198 | SNS-032 | CDK16;CDK17;CDK2;CDK7;CDK9;CDKL5 | inhibitor of cyclin-dependent kinases |
| 6.4544 | 8.4468 | 10.961 | vincristine | | inhibitor of mictrotubule assembly |
| 7.9384 | 8.4809 | 12.683 | pluripotin | MAPK1;RASAL1 | promoter of embryonic stem cell self-renewal; inhibitor of Ras-GAP and ERK |
| 8.1199 | 8.5169 | 10.448 | tanespimycin | HSP90AA1 | inhibitor of HSP90 |
| | 8.9724 | 14.457 | bleomycin A2 | | inducer of DNA damage |

| | | | | | |
|---|---|---|---|---|---|
| | 8.9939 | 9.1898 | bardoxolone methyl | | electrophilic inducer of the NFE2L2-KEAP1 pathway |
| 3.0442 | 5.1868 | 6.64 | leptomycin B | XPO1 | inhibitor of exportin 1 |
| 4.7827 | 7.1199 | 6.7813 | LBH-589 | HDAC1;HDAC2;HDAC3;HDAC6;HDAC8 | inhibitor of HDAC1, HDAC2, HDAC3, HDAC6, and HDAC8 |
| 5.08 | 5.4913 | 7.217 | ouabain | ATP1A1;ATP1A2;ATP1A3;ATP1A4;ATP1B1;ATP1B2;ATP1B3;ATP1B4 | cardiac glycoside; inhibitor of the Na+/K+-ATPase |
| 10.709 | 12.088 | 8.3445 | CAY10618 | NAMPT | inhibitor of nicotinamide phosphoribosyltransferase |
| 6.5894 | 10.441 | 8.8831 | dasatinib | EPHA2;KIT;LCK;SRC;YES1 | inhibitor of SRC, YES1, EPHA2, c-KIT, and LCK |
| 9.4129 | 9.9905 | 8.9214 | AZD7762 | CHEK1;CHEK2 | inhibitor of checkpoint kinases 1 and 2 |
| | 8.9939 | 9.1898 | bardoxolone methyl | | electrophilic inducer of the NFE2L2-KEAP1 pathway |
| 11.353 | 10.33 | 9.2588 | SCH-79797 | F2R | antagonist of proteinase-activated receptor 1 (PAR1) |
| | 4.6395 | 9.3541 | CR-1-31B | EIF4A2;EIF4E;EIF4G1 | silvestrol analog; inhibits translation by modulating the eIF4F complex |
| 14.43 | 13.901 | 9.5437 | epigallocatechin-3-monogallate | | natural product |
| 7.4058 | 8.3521 | 9.7719 | MLN2238 | PSMB5 | inhibitor of 20S proteasome at the chymotrypsin-like proteolytic (beta-5) site |
| 9.4783 | 12.377 | 9.8555 | bosutinib | ABL1;SRC | inhibitor of SRC and ABL1 |
| 7.7222 | 7.3893 | 9.9091 | SR-II-138A | EIF4A2;EIF4E;EIF4G1 | silvestrol analog; inhibits translation by modulating the eIF4F complex |
| 6.7611 | 6.8773 | 10.028 | omacetaxine mepesuccinate | | inhibitor of protein translation by preventing protein elongation |
| 1.9382 | 9.7431 | 10.082 | oligomycin A | ATP5L2 | inhibitor of mitochondrial ATP synthase |
| 10.201 | 11.038 | 10.093 | obatoclax | BCL2;BCL2L1;MCL1 | inhibitor of MCL1, BCL2, and BCL-xL |
| 7.012 | | 10.157 | canertinib | EGFR;ERBB2 | inhibitor of EGFR and HER2 |
| 6.3105 | 9.1778 | 10.217 | 1S,3R-RSL-3 | GPX4 | synthetic lethal with HRAS in engineered cells; inhibitor of GPX4 |
| 7.8567 | 9.2225 | 10.323 | AZD8055 | MTOR | inhibitor of mTOR |
| 5.8898 | 9.0936 | 10.362 | afatinib | EGFR;ERBB2 | inhibitor of EGFR and HER2 |

[1]Data from: Seashore-Ludlow, Brinton, et al. "Harnessing connectivity in a large-scale small-molecule sensitivity dataset." *Cancer discovery* 5.11 (2015): 1210-1223. DOI: 10.1158/2159-8290.

[2]Values for each cell line represent area-under-concentration-response curve (AUC, percent viability plotted as a function of inhibitor concentration) for each compound–cancer cell line pair, as a measure of cell sensitivity to small-molecule treatment. Smaller values represent greater sensitivity.

**Supplementary Table 5.** Mutations shared between more- and less-responsive cell lines in our experiments

| More responsive (SCaBER and SW780) | Less responsive (RT4 and HT-1376) |
|---|---|
| *NGEF* | *TTN*[2] |
| *PAPPA2* | *MUC4* |
| *RPGRIP1L* | *DCHS2* |
| *HYDIN* | *MAMDC4* |
| *ABR* | *PHLDA1* |
| *SIN3B* | *GNPTAB* |
| *TTN*[2] | *MARK3* |
| *ABCA12* | *LOXHD1* |
| *SLC5A4* | *SNAPC4* |
| *DDX60L* | *NBPF15* |
| *EGFLAM* | *KIF1A* |
| *DNAH8* | |
| *SASH1* | |
| *OPLAH* | |
| *MT-CO1* | |
| *MT-CO3* | |
| *MT-ND4* | |
| *MT-ND5* | |
| *MDN1* | |

[1]Data from: The Cancer Cell Line Encyclopedia (https://portals.broadinstitute.org/ccle/about)
[2]This was the only gene shared between the two pairs

**Supplementary Figure 1.** Frequency of gene mutations shared between more- and less-responsive cell lines in our experiment, in bladder cancer patients. Frequency shown is in human bladder cancer patients. Data from The Cancer Genome Atlas (https://doi-org.prox.lib.ncsu.edu/10.1016/j.cell.2017.09.007)

**A. Mutations observed in sensitive cells in our study (SCaBER and SW780)**

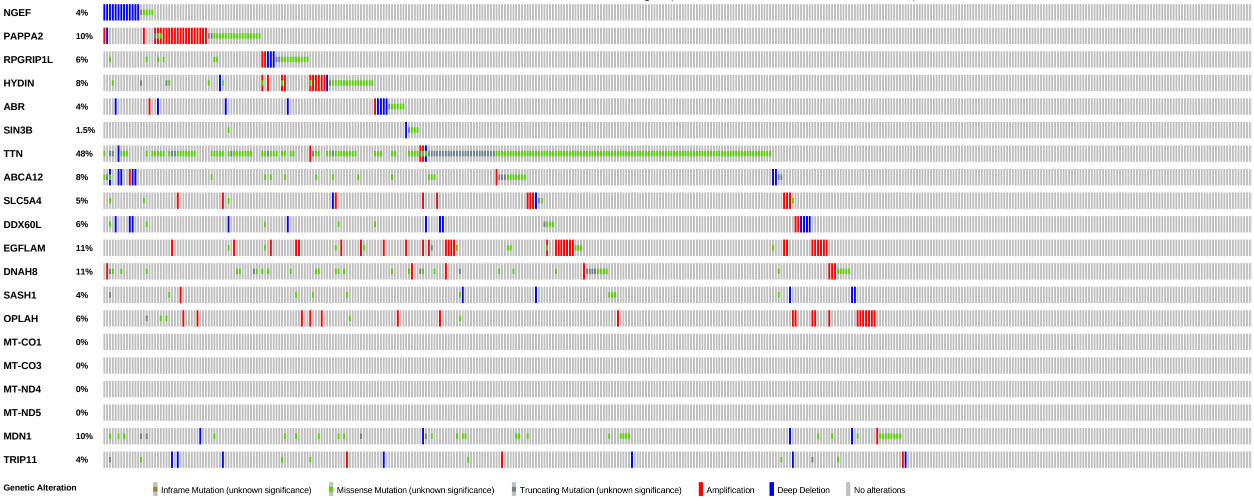


**B. Mutations observed in resistant cells in our study (RT4 and HT-1376)**

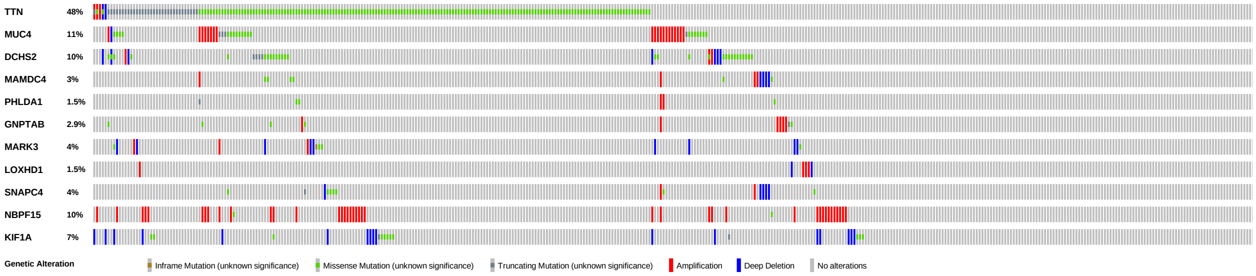
TTN 48%
MUC4 11%
DCHS2 10%
MAMDC4 3%
PHLDA1 1.5%
GNPTAB 2.9%
MARK3 4%
LOXHD1 1.5%
SNAPC4 4%
NBPF15 10%
KIF1A 7%
Genetic Alteration
Inframe Mutation (unknown significance)
Missense Mutation (unknown significance)
Truncating Mutation (unknown significance)
Amplification
Deep Deletion
No alterations
